# Deciphering antigen-driven T cell responses through vectorized TCRdist sequence neighborhood quantification

**DOI:** 10.64898/2026.04.10.717405

**Authors:** Sebastiaan Valkiers, Koshlan Mayer-Blackwell, Albert C. Yeh, Vincent Van Deuren, Andrew Fiore-Gartland, Geoffrey R. Hill, Kris Laukens, Pieter Meysman, Philip Bradley

**Author notes:** These authors contributed equally (Pieter Meysman, Philip Bradley).

## Abstract

T cells provide precise mechanisms to defend the body against infection and malignancies, mediated through the expression of their hypervariable T cell receptors (TCRs). Interpreting similarity between TCRs, however, remains a significant challenge. While performant clustering methods exist, these often fail to distinguish between antigen-driven convergent selection and patterns arising stochastically from biases in the V(D)J recombination mechanism. Moreover, defining enrichment in sequence similarity among large repertoires is computationally taxing. To address these limitations, we present an efficient computational framework for rapid approximation of TCRdist distances using fixed-length vector embeddings and highly optimized nearest neighbor search, allowing sequence similarity enrichment testing at a multi-repertoire-wide scale. This framework leverages a novel shuffling-based background model that preserves important repertoire characteristics such as V gene frequency, CDR3 sequence length and generation probability more accurately than synthetic models. Together, these tools enable the efficient and robust identification of significantly neighbor enriched (SNE) TCR sequences at scale. We validate this approach by showing a significant enrichment of SNE clones in memory T cell fractions and further demonstrate its utility in identifying convergent T cell signatures of response to vaccination and viral infections, providing a scalable approach for antigen-agnostic T cell response profiling.

## Introduction

T-cell receptor (TCR) repertoires are highly diverse and individualized due to allelic diversity, HLA polymorphism, and quasi-random recombination of germline encoded V, D and J gene segments. Differential selection and expansion events resulting from exposure to specific antigens further drive the unique composition of the TCR repertoire in each individual. Acutely expanded T cell clones may undergo a transition towards a memory phenotype, providing long-term protection against infection. These transitions are characterized by a contraction in clonal frequency of previously expanded clones. These clones remain nonetheless detectable, thereby leaving an epitope-specific signature in the repertoire. A critical task is the identification of those biologically meaningful signals of recent antigen exposure within complex TCR repertoires. Recognition that receptors with shared epitope specificity display higher-than-average TCR sequence similarity provides a basis for exploring the TCR landscape for antigen-mediated convergent selection. However, clonal convergence is driven by two main processes: convergent recombination and/or convergent selection. Since the V(D)J recombination process is biased, certain sequences or motifs may be preferentially generated during development and present in repertoire samples without antigen-mediated selection [1]. This recombination bias primarily occurs due to the selection of germline genes, alongside factors such as the quantity of deletions and the insertion of non-templated nucleotides during TCR formation.

Several statistical formulations attempt to capture the recombination bias [1–3]. These models permit the estimation of the probability of independent gene recombination events, also known as a receptor’s probability of generation (P_gen_). Yet, due to the dynamic and responsive nature of the adaptive immune system, clonal distributions of empirical samples differ substantially from the clonal distributions that are generated by these models. These differences are most plausibly driven by selection mechanisms imposed by the interaction of HLA molecules and self-proteins during negative and positive thymic selection during development [4] and later exposure to non-self antigens throughout life. Clonal selection theory posits that T cells expressing receptors with sufficient affinity for HLA-peptide ligands proliferate during an immune response. Since similar TCRs are more likely to have similar affinity for a particular HLA-peptide ligand, such antigen-driven selection may result in a detection of groups of T cell clones with high sequence similarity [5,6]. For example, Pogorelyy and colleagues (2019) illustrated that clones with larger than expected sequence similarity overlap with antigen-responding clones from single repertoire snapshots [7]. While prior studies have proposed evaluating enrichment in clones with shared sequence similarity–relative to statistical models of receptor probability of generation–more computationally efficient approaches are needed to perform sequence neighbor density estimation in large immune receptor datasets. Moreover, most sequence similarity measurements have relied on discrete thresholds, such as an edit distance of 1, which may limit the resolution needed to capture more nuanced relationships between immune receptor sequences.

To effectively discriminate between clusters of similar TCRs arising from antigen-mediated selection versus recombination biases, we propose a framework to identify statistically anomalous convergent recombination events. To implement this framework, we embed the complementarity-determining regions (CDRs) of the TCR into a fixed length vector encoding, permitting an ultra efficient euclidean distance based approximation of the TCRdist metric [5,8] (vecTCRdist) compatible with highly optimized nearest neighbor search libraries such as faiss [9]. We further propose a shuffling-based method for estimating each clone’s expected neighbor frequency by chance alone. Combining these approaches, we show that vectorized TCRdist enables computationally efficient neighborhood enrichment testing on a continuous distance scale, and is applicable to both single-chain TCRα/β or paired-chain TCRαβ datasets. We find that significantly neighbor-enriched (SNE) clones are a hallmark of response to an attenuated viral vaccine and an imprint of common viral infections, suggesting that our framework has broad potential implications for identifying T cell signatures of immunological control.

## Methods

### Datasets

#### Public data

We selected a range of publicly available human TCR data sets that have been widely used in the field of AIRR data analysis for method development and benchmarking [10-15]. The different public data sets used in this study are listed in Table S1.

#### Murine naive and memory T cells

In addition, we generated a new dataset of mouse TCR sequences from sorted naive and memory T cells to evaluate model performance in a second organism. All mice were housed in the animal facility at the Fred Hutchinson Cancer Center, kept in sterilized microisolator cages, and received acidified autoclaved water (pH 2.5). Splenocytes were obtained following 70mm filtration of mechanically ground spleen and treatment with Gey’s solution for RBC lysis. Naive and memory CD4 T cells were flow sorted from a pool of 15 C57BL/6J mouse spleens. (naive: CD3+/CD4+/CD8-/CD62L+/CD44-; memory: CD3+/CD4+/CD8-/CD62L-/CD44+). Antibodies clones used include CD3 (PerCP/Cy5.5, Clone 17A2), CD4 (PacBlue, Clone RM4-5), CD8 (BV510, Clone 53-6.7), CD44 (APC/Cy7, Clone IM7), CD62L (PE, Clone MEL14). Genomic DNA extraction from flow-sorted cells was performed using the QIAamp DNA Mini Kit (QIAGEN, Cat#51304) as described in the product protocol, and TCR sequencing was performed using the mouse TCRB assay (v3) by Adaptive Biotechnologies.

### TCRDist-based encoding

TCRdist is a distance metric that takes into account the weighted dissimilarity of the amino acid sequences between any two TCRs [5,8]. TCRdist uses amino acid sequence information from four pMHC-contacting loops in the TCR, known as complementarity determining regions (CDRs). Three loops originate from the V gene portion of the TCR (CDR1, CDR2 and CDR2.5). The hypervariable CDR3 region describes the fourth loop. Loop definitions for CDR1, CDR2 and CDR3 are based on the IMGT CDR definitions (https://www.imgt.org/IMGTScientificChart/Nomenclature/IMGT-FRCDRdefinition.html). In addition, CDR2.5 is defined by the residues of the IMGT alignment spanning position 81-86 of the V domain.

In the present study we engineered a numeric encoding to closely approximate TCRdist, permitting the transformation of TCR receptors sequences of any length into a fixed-length numeric vector encoding, whose euclidean distances reflect the TCRdist between the underlying V gene and CDR3 (**Figure 1A**). First, the CDR3 region is trimmed at the 3rd and 2nd to final position. Following trimming, gaps are introduced at a fixed position to standardize the length of each sequence. Every amino acid (including the gap character) is mapped to its corresponding numerical vector. These amino acid vectors are concatenated in the same order as they appear in the sequence. To obtain an encoding for each amino acid, we use the rows of the 21x21 TCRdist distance matrix, where each row and column represents an amino acid. The TCRdist distance matrix is created by transforming the BLOSUM62 substitution matrix as follows: *dist(a,a)*=0; *dist(a,b)*=min(4, 4-BLOSUM62(a,b)). An additional row with a gap character is introduced. Substituting any amino acid with a gap character results in a gap penalty of 4. We take the square root of the TCRdist distance matrix to satisfy the triangle inequality. To decrease the total size of the final embedding, we use multidimensional scaling (MDS) to project the rows of the TCRdist distance matrix into an *n*-dimensional embedding, creating a dictionary of amino acid vectors. The resulting numerical representation captures the relative distance properties of each amino acid. Throughout the rest of this manuscript, this encoding will be referred to as the *vecTCRdist* embedding.

**Figure 1.**
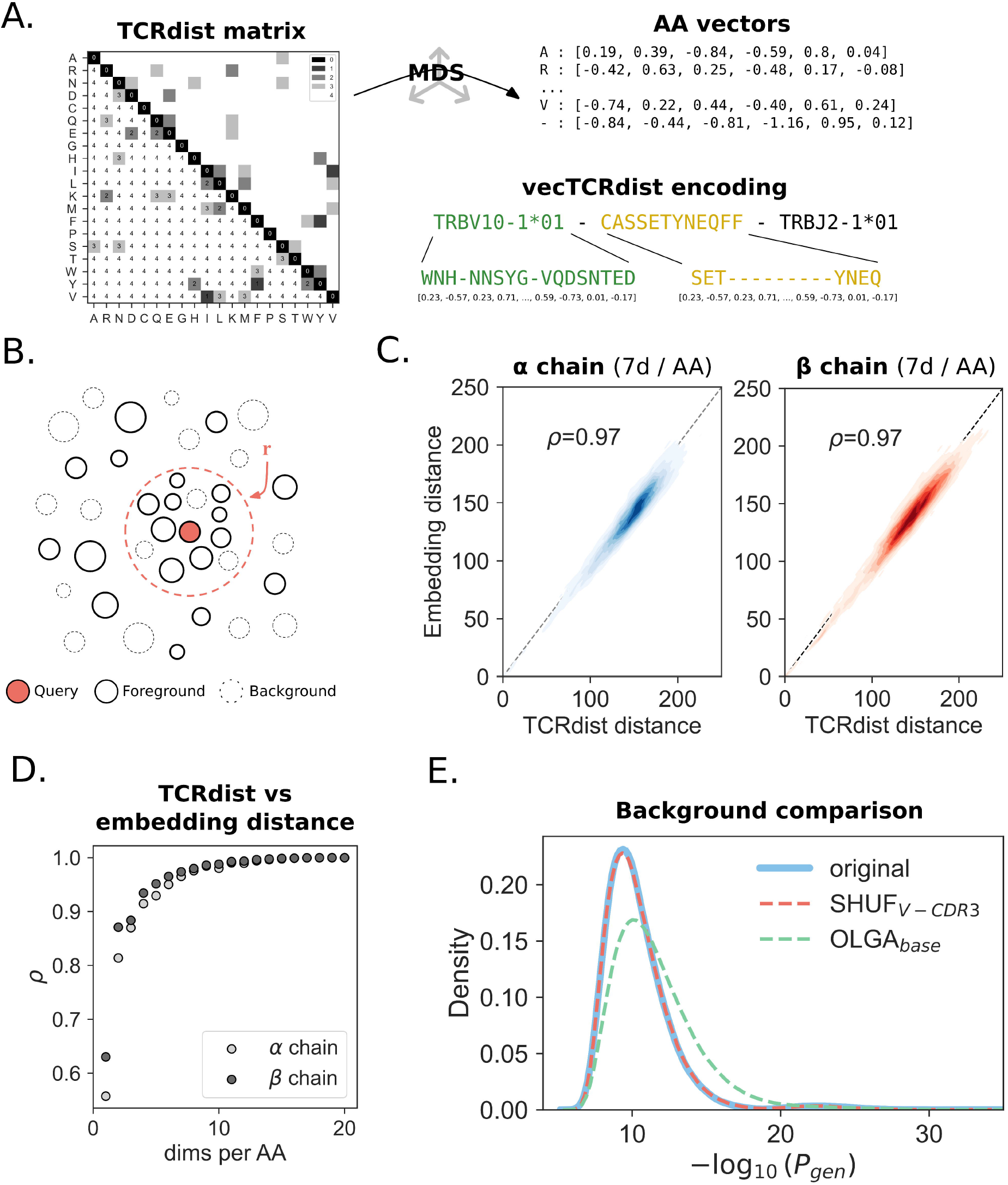
Using vecTCRdist distances to identify significantly neighbor enriched (SNE) TCRs. **A**. TCRdist distance information is projected into a lower dimensional vector representation using multidimensional scaling. Amino acid vectors are then concatenated into a vectorized representation for the full TCR. TCRdist distances are approximated by computing the squared euclidean distance between vectors. **B**. Any TCR in the dataset (reference TCRs) that falls within the radius r of the query TCR belongs to that TCR’s neighborhood. As such, the TCR neighborhood differs from a TCR cluster in that it is unique for each individual TCR. **C**. Comparison of squared euclidean distance between vecTCRdist embeddings and TCRdist distance for 924 αβTCRs. The analysis shows a near-perfect approximation of TCRdist distance using an encoding scheme that includes 7 dimensions per individual amino acid, resulting in a total embedding size of 259. **D**. Pearson correlation (*ρ*) between the squared euclidean distance of vecTCRdist embeddings and TCRdist distance for 924 TCRα and TCRβ sequences. Different embedding lengths were created by changing the number of dimensions per amino acid encoding in the MDS transformation. **E**. Generation probability distribution of background repertoires generated using the OLGA_base_ and SHUF_V-CDR3_, compared to the original input repertoire.

### Validating the vecTCRdist embedding

We used a set of 924 unique αβTCRs from the *COVID cross-reactivity* dataset to evaluate the correlation between the vecTCRdist embeddings and the TCRdist distances. α and β chains were encoded separately and pairwise Euclidean distances were calculated among the two chains. Similarly, we calculated the TCRdist between each pair separately for α and β using the tcrdist3 package [8]. Correlation between euclidean vecTCRdist and TCRdist distances was determined using the Pearson correlation coefficient.

### Similarity search

We use the faiss [9] library to store TCR vectors on an index, which allows rapid retrieval of distances between any two vectors. The vecTCRdist framework supports the use of both exact and approximate index structures, allowing a trade-off between computational performance and accuracy. A radius search can be performed on the index in order to retrieve all vectors that are within a Euclidean distance *r* from the query vector (**Figure 1B**). We refer to the set of all TCRs within *r* as the TCR neighborhood. By default, *r* was set to 12.5 for single-chain analyses. We used a set of repertoires with different sizes (ranging from 10,000 to 500,000 unique clones) to evaluate the computational performance of our method.

### Background

#### Synthetic background

We use the Optimized Likelihood estimate of immunoGlobulin Amino-acid sequences (OLGA) model described in Sethna et al (2019) [3] to generate *de novo* TCR sequences. We use the base P_gen_ model as the baseline for our background comparisons (OLGA_base_). In addition, we modified the OLGA model to select TCRs with a specific V gene. This enabled us to match the V gene frequencies in our samples through selective generation of TCRs. In addition, we aimed to match the frequency distribution for the lengths of the CDR3 amino acid sequence in the input repertoire. After determining the CDR3 length distribution we sample sequences with a specific V gene using an accept-reject strategy to match the sequence length distribution for every V gene in the input repertoire. This strategy results in a set of background TCRs that match the properties of the input repertoire in terms of V gene frequencies and CDR3 sequence length distribution. Hereafter, this background model is referred to as the OLGA_V-CDR3_ model.

#### Shuffled background

The shuffled background model attempts to generate a novel set of TCR sequences by resampling from an original repertoire, thereby preserving important properties such as germline gene frequencies and length distribution of the CDR3 amino acid sequences. First, breakpoints are established by identifying the V(D)J junctions from the CDR3 nucleotide sequence. During this process, each part of the nucleotide sequence is assigned to originate from the V, D or J germline gene segment, or from an N-insertion event. The breakpoint can only occur at the junctions, i.e. in between the V, D and J segments. Sequences are split at the breakpoint and subsequently shuffled. Valid breakpoints are located at the junctions between non-germline (N) insertions and germline sequences (V, D and J gene segments). The location of the breakpoints are critical for ensuring that the ‘recombination’ between two different TCRs is biologically feasible. The segments are randomly paired and joined if their breakpoints are compatible; that is, if they both have a breakpoint within the same type of region (pre-D or post-D) in their nucleotide sequences. During this process, new TCRs are accepted in a way that preserves the overall length and distribution of n-inserted nucleotides. As a result, the background set of TCRs will follow the same P_gen_ distribution as the original repertoire. Hereafter, this background model is referred to as the SHUF_V-CDR3_ model.

#### Background depth

We evaluated the influence of background depth by comparing TCR neighbor retrieval (*TCRdist*<12.5) in various background sizes. The background size is always calculated relative to the size of the input repertoire. We evaluated the following background ratios: 1:1, 1:2, 1:5, 1:10, 1:25, 1:50, 1:100, 1:250, and 1:500. We used the largest background (ratio 1:500) as a benchmark reference for accurate background neighbor estimates.

### Defining neighbor enrichment

In order to quantify the significance of the neighborhood size for a certain clonotype, we evaluate the likelihood of observing at least *k* neighbors using the hypergeometric distribution.

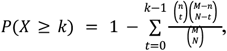

where *M* is the size of the total population (sample and background), *N* is the size of the sample of interest, *n* represents the total number of neighbors in the population and *k* describes the number of neighbors in the sample. To compute the probability of observing at least *k* neighbors *P*(*X* ≥ *k*), we use the survival function, which equates to the inverse sum of observing (0, 1,…, *k* − 1) neighbors. To limit the family-wise error rate in our results, Bonferroni correction was applied to adjust the p-values of the hypergeometric tests to account for the number of tests (i.e., the number of foreground TCRs). Hereafter, the method for determining TCR neighborhoods will be designated clustcrdist.

### Epitope-specificity annotation

TCR clonotypes were annotated using the VDJdb (version June 13, 2024) [16]. In order to limit the inclusion of likely false positive identifications, we excluded one particular study from the VDJdb with the following reference ID: https://www.10xgenomics.com/resources/application-notes/a-new-way-of-exploring-immunity-linking-highly-multiplexed-antigen-recognition-to-immune-repertoire-and-phenotype/. A flexible matching criterion was used, defining matches within a <24 TCRdist distance threshold for single chain data and <90 for paired chain TCRs, to connect similar TCRs. We then used the Leiden algorithm to identify communities in the graph, separating out the clusters.

### Hardware and software environment

All benchmarking experiments were performed on a 64-bit machine with 15.3 GB RAM, and an Intel® Core™ i7-10875H CPU @ 2.30GHz (8 cores, 16 threads), running Ubuntu 20.04.2.LTS. The machine also featured 256 KiB of L1 cache, 2 MiB of L2 cache, and 16 MiB of L3 cache.

The software used for developing the encoding strategy and TCR neighbor enrichment method was written in Python (3.9). Statistical analyses were conducted using the scipy (1.8.0), statsmodels (0.14.0), and scikit-learn (1.2.0) packages. Similarity searches were performed with the faiss (1.7.2) package. For data visualization, matplotlib (3.6.2) and seaborn (0.12.2) were utilized.

## Results

### Euclidean distances between TCRdist vectors reflect TCRdist distances

A fixed length representation is created for each TCR by trimming and gapping the CDRs (1, 2, 2.5 and 3). Next, each amino acid or gap character is mapped to a vector, which is created by scaling the TCRdist distance matrix into different dimensions ranging from 2 to 20 using MDS (**Figure 1A**). To test whether Euclidean distances between vectorized embedding recapitulated the TCRdist metric, we selected 924 TCRs for which full length alpha and beta chain information was available. For each TCRa and TCRb separately, an embedding was created. We next determined all pairwise TCRdist distances between the sequences, as well as all the (squared) euclidean distances between all embeddings. We found that high correlation between the two methods can already be achieved with as little as 7 dimensions (Pearson correlations of 0.97 and greater, **Figure 1C,D**).

Next, we tested the performance of the method in retrieving neighbor counts. We observed a linear increase in runtime as the number of sequences increased during index construction. In contrast, searching the index scaled quadratically. We then evaluated the computational performance of clustcrdist in determining pairwise TCRdist distances between paired αβ TCRs. This analysis may be run in one of two modes: exact (using the flat index) or approximate (using the ‘inverted file’ or IVF index). We compared the results of the IVF index with the flat index to assess the extent of accuracy loss with the approximate (IVF) method. To do this, we selected 200,000 αβTCRs and computed the pairwise TCRdist distances between them under a maximum threshold of 96.5 vectorized TCRdist units. The number of dimensions per amino acid was 16. We identified a total of 185,205 unique neighbor edges under the distance threshold using the exact method. The approximate method was able to capture 99.2% (183,752) of these edges (**Figure S1A**) at approximately 7x the speed (4301±577 vs. 599±144 seconds for flat and IVF index respectively) of the exact method (**Figure S1B**), suggesting the use of the IVF index is acceptable for larger datasets where computational time prohibits exhaustive search.

### Background model properties

In order to quantify neighborhood enrichment, the need for an adequate null distribution is essential. We explored properties of different synthetic TCR backgrounds, using 3 different strategies to generate TCR sequences in a 10:1 ratio relative to the repertoire of interest: OLGA_base,_ OLGA_RS_ and SHUF_V-CDR3_. OLGA_base_ uses the OLGA implementation of the IGOR method to generate sequences. In OLGA_RS_, we modified OLGA_base_ with a rejection-sampling approach to match the V gene usage frequency as well as the CDR3 amino acid length distribution (OLGA_RS_). Finally, the SHUF_V-CDR3_ method uses sequences from the input repertoire and reshuffles them to generate a set of background TCRs. We first compared the repertoire properties of the OLGA_base_ and SHUF_V-CDR3_ models. For the OLGA_base_ model, it was found that the V gene usage frequency differed drastically from the input repertoire (**Figure S2A**). In contrast, this property was preserved with the SHUF_V-CDR3_ model, where the V gene usage matched up near-perfectly. When subsequently looking at the CDR3 length distribution, this was clearly shifted to the right in the OLGA_base_ background set (**Figure S2B**), meaning that OLGA_base_ generates TCRs with comparatively longer CDR3 amino acid sequences compared to the input repertoire. The SHUF_V-CDR3_ model, on the other hand, was able to model the CDR3 length distribution with high accuracy. As a result of the shifted CDR3 length distribution in OLGA_base_, the TCRs resulting from this model have a lower P_gen_ compared to the reference distribution (**Figure 1E**). The sequences resulting from SHUF_V-CDR3_, on the other hand, follow a nearly identical P_gen_ distribution as the TCRs in the input repertoire (**Figure 1E**).

Another important consideration is the size of the background. It was hypothesized that larger backgrounds would provide more accurate estimates of the expected neighbor distribution of the reference sample. Therefore, this parameter would have a substantial influence on the results of any TCR neighborhood enrichment analysis. The influence of background depth was evaluated by comparing neighbor retrieval (*r*<12.5) in various background sizes for TCRβ sequences. The background size was always calculated relative to the size of the input repertoire. We evaluated the following background ratios for 4 different samples in the *YFV cohort*: 1:1, 2:1, 5:1, 10:1, 25:1, 50:1, 100:1, and 250:1. The largest background (ratio 1:250) was used as a benchmark reference for accurate background neighbor estimates. For one of the samples, an additional 500x background was generated as an extreme case to study how closely this would reflect the neighbor distribution of the reference sample. First, we studied the neighbor distribution obtained at different background depths. It was found that any background datasets with >10:1 ratio to the input data showed minimal improvement in terms of matching the original neighbor distribution (**Figure S3A**). Next, we investigated whether increasing the size of the background data would produce different results in terms of the total number of SNE TCRs. As a reference, we compared against the deepest background (250:1). There was a sigmoidal relationship between the size of the background dataset and the number of neighbor-enriched sequences. For all samples the transition between the 5:1 to 10:1 background marked the transition from exponential to plateau phase of the curve (**Figure S3B**). When compared to the 250:1 reference, there is a significant increase in both the number of identified SNEs between the 2:1 and 5:1 background (**Figure S3B**) and their overlap with the SNEs identified using the reference background (**Figure S3C**). Based on the above results, it is reasonable to conclude that, using a background with a 10:1 size ratio to the input repertoire is a good strategy for capturing the most highly neighbor-enriched TCR sequences, while balancing computational efficiency and manageable search times.

Thirdly, we investigated how the different background models estimate neighbor density – minimizing sample-specific biases from background models introduced by differences in V gene usage and P_gen_. The data generating process for simulating a null background strongly influences the statistical significance of the observed neighbor counts in the repertoire. If the background does not accurately reflect the input repertoire, this may lead to over-or under-estimation of the neighbor enrichment. Consistent with previous research [7], we observed that an OLGA-generated background grossly underestimates the neighbor density (**Figure S4A**,**B**). Even after correcting for V gene usage and CDR3 length, the OLGA_V-CDR3_ method underestimated the empirically observed neighbor density by a factor 10 (**Figure S4A**,**B**), meaning that the number of estimated neighbors by the model is tenfold smaller than the number of observed neighbors in the repertoire. One way to account for the underestimated neighbor count, is through the application of a selection factor *q* [7]. The neighbor distribution estimated by the SHUF_V-CDR3_ method, on the other hand, correlated strongly with the observed neighbor distribution in the repertoire, and thus the use of a selection factor was not necessary in this case. Sequence generation models that attempt to account for thymic selection (e.g, SONIA [4]) may not account for individual factors (for example, V and J gene usage biases) or amplification/sequencing modality that may affect background distribution. Collectively, these results favor the SHUF_V-CDR3_ method as the conservative method of estimating expected neighbor frequency in absence of antigen-mediated selection.

### Validating the SNE TCR identification framework

TCR encoding permits rapid computation of vectorized TCR distances between all members of a repertoire and radius search, which we leverage to compute the neighbor distribution of each clone. By comparing the observed distribution of neighbor counts with a simulated background neighbor distribution, we sought to identify anomalies potentially indicative of antigen-driven selection. We refer to sequences with more neighbors than would be expected by chance (*p*, Bonferroni <0.05) as significantly neighbor enriched (SNE) clones. Since the SHUF_V-CDR3_ method for background generation makes use of the sequence properties present in the repertoire, we observed no need for the application of a correction factor that accounts for thymic selection (**Figure S4A**,**B**). Moreover, the use of the SHUF_V-CDR3_ model revealed two neighbor-enriched populations that were not detected using either OLGA-based backgrounds (**Figure S4C**,**D**), suggesting the potential utility of the SHUF_V-CDR3_ model to more sensitively identify antigen-driven selection within TCR repertoires.

To validate our approach, we started by exploring the distinct characteristics of naive and memory T cells. Naive T cells have not yet experienced strong antigen selection that would be expected to drive strong polyclonal expansion of similar receptors and give rise to SNE clones. Thus, we hypothesized that a comparison of naive and memory T cell repertoires might reveal a differential number of SNEs. We applied our neighborhood enrichment framework to a dataset of murine sorted naive and memory T cell fractions. As hypothesized, the memory T cell fractions contained significantly higher numbers of absolute SNE clone counts compared to the naive T cells (*p*=0.009, Mann-Whitney U test) (**Figure 2A**). Next, we examined human samples where T cells were sorted into naive, effector memory (EM) and central memory (CM) fractions (*n*=43, 21, 22 respectively). TCR neighborhood enrichment was determined for each sample. Again, significant differences were observed in the fractional SNE clone count between naive and both EM (*p*=0.05) and CM (*p*=0.01), but not amongst memory fractions (*p*=0.30) (**Figure 2B**). These results demonstrate how our workflow successfully identifies non-random instances of sequence similarity within the repertoires of both mice and humans.

**Figure 2.**
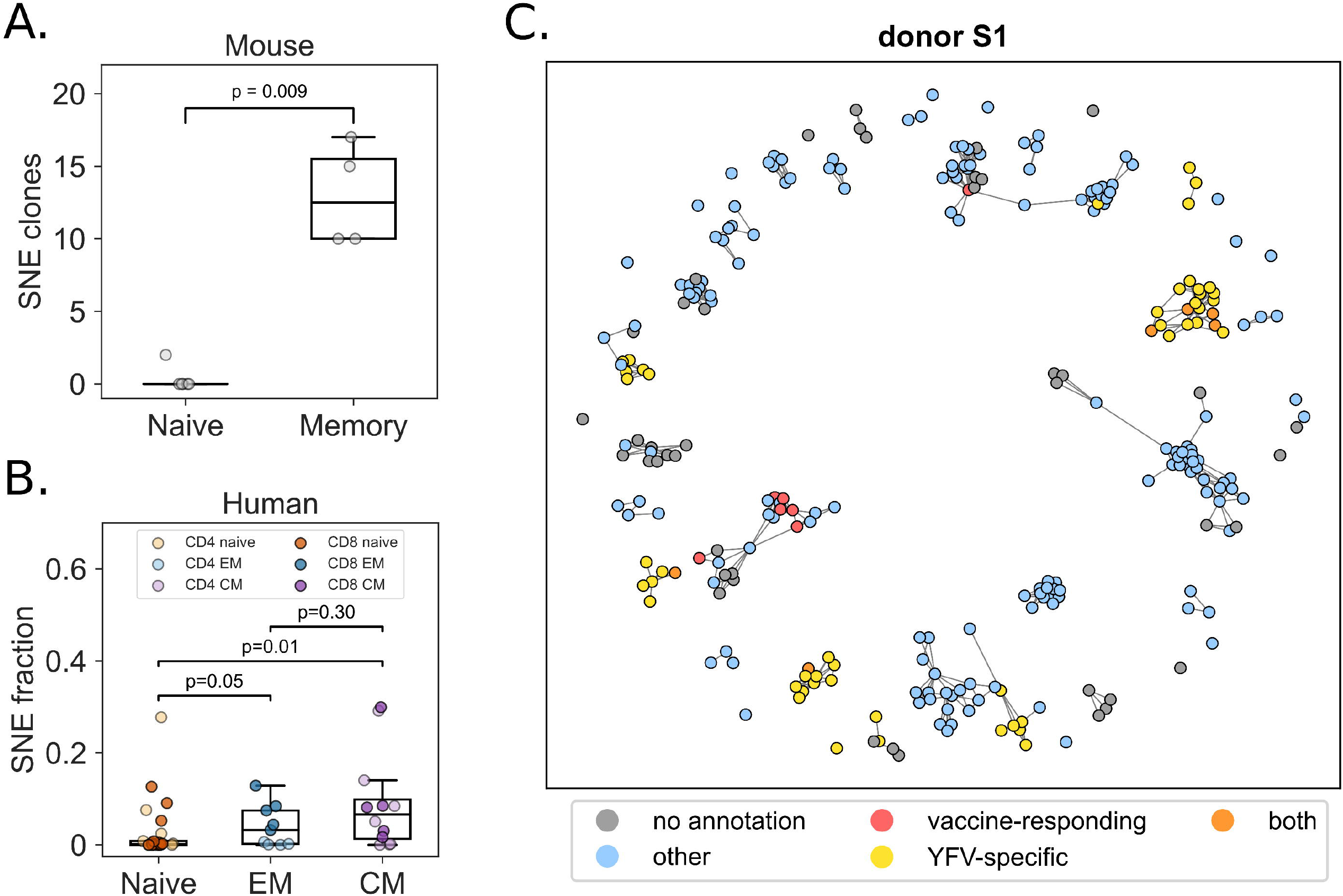
Validating the neighbor enrichment framework. **A**. The memory fraction of T cells extracted from mice display higher rates of clonal convergence compared to naive T cells from the same donor (p = 0.009). **B**. Similar results are observed for sorted human T cells. Higher SNE counts were measured in both effector memory (*p*=0.004) and central memory (*p*=0.007) fractions as compared to naive clones. **C**. Annotating SNE clones in a vaccination case study reveals convergent responses in YFV-specific T cell clones, and not necessarily in dominant clones that undergo longitudinal expansion. Red, vaccine-responding: clones that longitudinally expanded upon vaccination with the YFV vaccine; Yellow, YFV-specific: clones matching (*TCRdist*<24) with a YFV epitope-specific TCR in VDJdb; Orange, both: vaccine-responding clones that also have a YFV-specific match in VDJdb. Blue, other: clones matching (*TCRdist*<24) any epitope-specific TCR in VDJdb. Grey, no annotation: clones for which no annotation was found.

### Yellow fever virus vaccination induces convergent selection

We next applied the neighborhood enrichment strategy on a data set of 3 paired twins whose repertoires were sequenced at multiple timepoints after yellow fever virus vaccination [13]. In the original study by Pogorelyy and colleagues (2018), it was found that the T cell response to vaccination peaked at 15 days after injection. The authors curated a list of 5,969 vaccine-responding clones based on the expansion and subsequent contraction behavior. Here, we hypothesized to see higher levels of neighbor enrichment at the day 15 post vaccination timepoint, relative to the other measurements. Consistent with the validation experiments on mouse and human naive and memory T cell fractions, we performed our analysis on each repertoire individually. While we observed SNE clones across all timepoints, at day 15 post vaccination a larger proportion of the top enriched T cell clones originated from the pool of YFV vaccine-responding clones (**Figure S5A**). This observation potentially suggests that vaccination elicits convergent selection in the repertoire to some degree. Indeed, some of the longitudinally expanded clones (YFV vaccine-responding) saw an increase in neighborhood enrichment at the 15 day post-vaccination timepoint, which was absent at other timepoints (**Figure S5B**). However, for the majority of longitudinal expanders no neighbor expansion was observed (**Figure S5B**, red line indicates the average). Moreover, there was no clear relation between clone count and TCR neighborhood enrichment. Indeed, when we studied this relationship in more detail, we found no correlation between clone rank and neighbor enrichment p-value rank for any of the repertoires (**Figure S6**). We evaluated the overlap between the most abundant clones and those classified as SNE, finding it to be minimal, with an average of 0.5% (95% CI: 0–1.5%).

We then compared the results of our neighbor enrichment analysis with the previously curated list of vaccine-responding clones (in Pogorelyy et al., 2018 [13]). Interestingly, the neighbor enrichment response was more shared across individuals when compared to the vaccine-responding clones, independent of their specificity (**Figure S7A**). We observed that a large number of SNE clones were not part of the curated list of vaccine-responding clones identified in the original study, because they lacked sufficient longitudinal expansion. To investigate whether these SNE clones contributed to the vaccination response in another way than longitudinal expansion, we attempted to infer their epitope specificity. The strategy used for this was querying against the VDJdb, allowing a <24 TCRdist mismatch. We found substantial annotation with YFV epitopes at 15 days post YFV-vaccination (**Figure 2C, S7B, S7C, S8, Table S2**), suggesting that these SNE clones may indeed respond to the vaccination stimulus using the alternative mechanism of sequence convergence, rather than clonal expansion. Another set of SNE clones were found to have a high degree of specificity towards common viral epitopes including NLVPMVATV (cytomegalovirus, CMV), GLCTLVAML (Epstein-Barr virus, EBV) and GILGFVFTL (influenza A virus). Indeed, when looking at the annotation rate among SNE clones, compared to the vaccine-responding clones, SNEs display a higher specificity towards CMV (*p*=0.002) and EBV (*p*=0.002) epitopes, but not to YFV (*p*=0.180) (**Figure S7C**). To further investigate the role of specificity against viral epitopes, SNE clones were compared against the most expanded clones in the repertoire, specifically at the day 15 post-vaccination timepoint. For equal comparison, we took the *n* most expanded clones in each repertoire, where *n* is equal to the number of SNE clones identified in that repertoire. This comparison revealed an average 2-fold increase of YFV epitope-specificity annotation rate for the SNE clones compared to the top expanded clones (**Table S2**).

### Paired αβ chain data improves resolution of neighbor enrichment

Building on the YFV vaccination case study, where we examined only the TCRβ chain, we extended this approach by including both α and β chains in the current SARS-CoV-2 infection dataset. We used CD4/CD8-sorted paired αβ chain TCRs from a single donor which was longitudinally sampled during a SARS-CoV-2 infection – a baseline sample at 143 days pre-infection (“baseline”), as well as samples at 6 (“acute”) and 29 (“convalescent”) days post-infection. After filtering, the dataset contained a total of 102,554 TCRs (76,255 and 26,299 CD4^+^ and CD8^+^ respectively) across all timepoints. We were primarily interested in whether SARS-CoV-2 infection created a signature of convergent selection in the TCR repertoire at the acute timepoint. To this end, neighbor enrichment analysis was applied individually to each timepoint. We identified several stable SNE clusters in both the CD4^+^ and CD8^+^ fractions (**Figure S9A**,**B**). Notably, within the CD8^+^ population, a large cluster predominantly expressing TRAV1-2 (100%) with TRAJ33 (84%), as well as some TRAJ20 (9%) and TRAJ12 (6%), was enriched across all timepoints, likely representing a group of mucosal-associated invariant T (MAIT) cells (**Figure S9C**). Interestingly, even when excluding cells with MAIT characteristics, there were 5 times as many CD8^+^ clones uniquely identified as SNE compared to the CD4^+^ population (**Figure S9D**). The stable clusters in the CD4^+^ fraction were characterized by TRAV6*01 and a GTA motif in the CDR3α (cluster 0), and TRAV17*01/TRAJ39*01 (cluster 1) (**Figure S9E**).

Next, we sought to determine if any of the CD8^+^ SNE clusters could be linked to known epitopes by co-clustering the non-MAIT clusters with paired αβTCR chains in the VDJdb (**Figure 3A**). At baseline, none of the SNE clusters matched known epitope-specific TCRs. In the sample taken during acute infection, however, we identified 2 SARS-CoV-2-specific clusters, with sequences matching reference sequences annotated for reactivity with SARS-CoV-2 spike peptides HLA-B*07:02-restricted SPRWYFYYL and HLA-A*02:01-restricted YLQPRTFLL. Another SNE cluster matched reference receptors annotated as reactive to the HLA-A*02:01-restricted GLCTLVAML epitope from EBV. We observed a contraction in the frequency of clones of the SARS-CoV-2-specific clusters between the acute and convalescent timepoints. In contrast, the EBV-specific SNE clones were more stable. By integrating longitudinal relative abundance information, we examined whether the SNE clones overlapped with receptors that expanded between baseline and acute time points. We identified 2366 strongly expanded clones at the acute timepoint defined as those with a log_2_(fold change) of > 3 compared to the abundance in the baseline sample. Out of 1058 SNE clones, 322 (30%) were also among the acutely expanded (**Figure 3B**). Notably, while only 2.8% of longitudinally expanded clones co-clustered with SARS-CoV-2-specific TCRs in the VDJdb, this was true for 20% of SNE clones (excluding MAIT cells) (**Figure 3C**). Similarly, 0.9% of longitudinally expanded clones and 3.4% of SNE clones were associated with EBV-specific TCRs in the VDJdb (**Figure 3C**). When looking at clones that are both expanded and SNE, the proportions were even larger (25% and 5% for SARS-CoV-2 and EBV respectively). Finally, the MAIT cell cluster, while enriched, contained no significantly expanded clones at the acute timepoint (**Figure 3D**). Taken together, longitudinal expansion and neighbor enrichment appears to be the strongest indicator of T cell response to recent antigen exposure.

**Figure 3.**
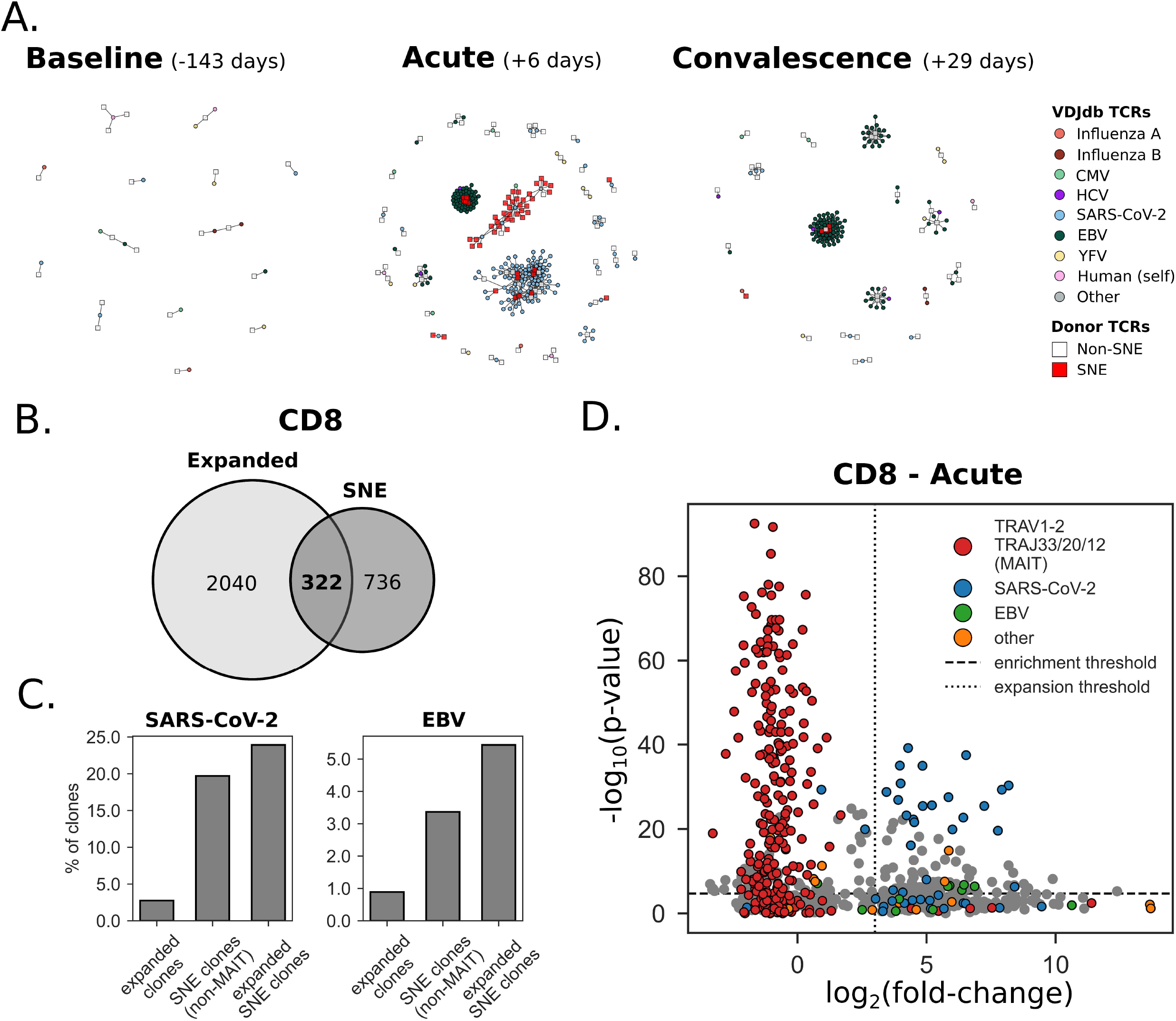
Neighbor enrichment analysis of paired αβTCRs from CD8^+^ T cells during SARS-CoV-2 infection. **A**. Co-clustering of donor CD8^+^ αβTCRs (>1 sequence neighbor) with TCRs in the VDJdb at baseline, acute infection and convalescence (left to right). The SNE clusters at the acute timepoint are enriched for SARS-CoV-2 and EBV epitopes. **B**. Intersection between longitudinally expanded (baseline vs. acute) and SNE clones. **C**. Percentage of clones co-clustering with SARS-CoV-2 or EBV-specific αβTCRs in VDJdb. **D**. Longitudinal expansion (as measured as the log_2_FC of the abundance at baseline vs. acute) at the acute phase of SARS-CoV-2 infection in relation to neighbor enrichment. MAIT cells are highly neighbor-enriched, but show no sign of longitudinal expansion. Among the longitudinally expanded SNE clones, ∼30% are associated with either SARS-CoV-2 or EBV.

### TCR neighborhood enrichment is dynamic across the lifespan

The *ages dataset* from a study by Britanova and colleagues contains TCR repertoire data from a unique cohort, encompassing blood samples from individuals across the full age spectrum, ranging from umbilical cord blood (UCB) to centenarians [12]. Here, we were interested to see if the degree of neighbor enrichment is variable across the lifespan. To this end, we used the *ages dataset*, containing a total of 77 unique samples from different individuals. To eliminate any biases related to repertoire size, repertoires were downsampled to 100,000 TCRs (based on abundance), thereby removing 8 subjects from the analysis whose repertoire contained < 100,000 unique clones. Subjects were categorized into different age groups, consistent to the definitions in the original study by Britanova et al. (2016). The age groups included UCB (*age*=0; *n*=8), young (*age*=6–25; *n*=12), middle age (*age*=30–50; *n*=11), aged (*age*=51–75; *n*=15), and long-lived (*age*=85–103; *n*=22). Neighbor enrichment analysis was performed on each (downsampled) repertoire. We next compared the number of SNE TCRs across different age groups and identified an increase from UCB to young and middle-age categories (*p*=0.05, Mann-Whitney U test, Benjamini-Hochberg correction) (**Figure S10A**). Interestingly, a decrease in SNE TCR count was observed in the aged and long-lived age groups, compared to individuals in the young and middle-aged group (*p*=0.02 and *p*=0.04 respectively, Mann-Whitney U test, Benjamini-Hochberg correction). Furthermore, when excluding the UCB samples, we observed a mildly decreasing tendency in overall neighbor enrichment with age (**Figure S10B**).

### SNE clones may be associated to (chronic) viral infections

In a previous experiment we identified that SNE clones potentially display a stronger tendency to bind common viral epitopes (e.g. from CMV and EBV). To further investigate this observation, we evaluated the binding specificities of SNE clonotypes in a large cohort containing samples from 666 individuals with known CMV serotype and HLA type [15]. First, neighbor enrichment analysis was performed on all repertoires. SNE clones were found in 583 out of 666 subjects. The SNE clones were annotated using a reference of known TCR-epitope interactions obtained from the VDJdb. An exact matching strategy was applied, requiring identity between the query and reference CDR3 amino acid sequence, V-gene, and J-gene. The annotation rate was subsequently compared across three common viruses: CMV, EBV, and Influenza A virus. These were selected based on the large number of known epitope-specific TCRs that are available for these species.

We compared the annotation rate of SNE clones to 3 control scenarios: (i) “top clones”, (ii) P_gen_-matched synthetic clones, and (iii) random clones. Top clones are those with the highest frequency in the repertoire regardless of neighbor enrichment score. The SHUF_V-CDR3_ model was used to generate the P_gen_-matched synthetic clones. We used OLGA to estimate P_gen_ of each receptor, confirming that the P_gen_ distribution of the SHUF_V-CDR3_ clones aligned with the P_gen_ distribution SNE clones. Compared to the control sets, SNE clones much more frequently matched reference TCRs with previously annotated epitope specificity (**Figure 4A**). Specifically, there were more CMV and Influenza A virus epitope-specific TCRs among SNE clones compared to the top clones (*p*=7.5e-07 and *p*=3.165e-21, respectively). Following this, the top clones also included a significant portion of epitope-specific TCRs, greater than the P_gen_-matched clones for all viruses. Top clones were more likely to match a reference sequence than randomly drawn clones, reflecting the fact that clonal abundance may encode some information about prior clonal expansion to common or recurrent immunological challenges.

**Figure 4.**
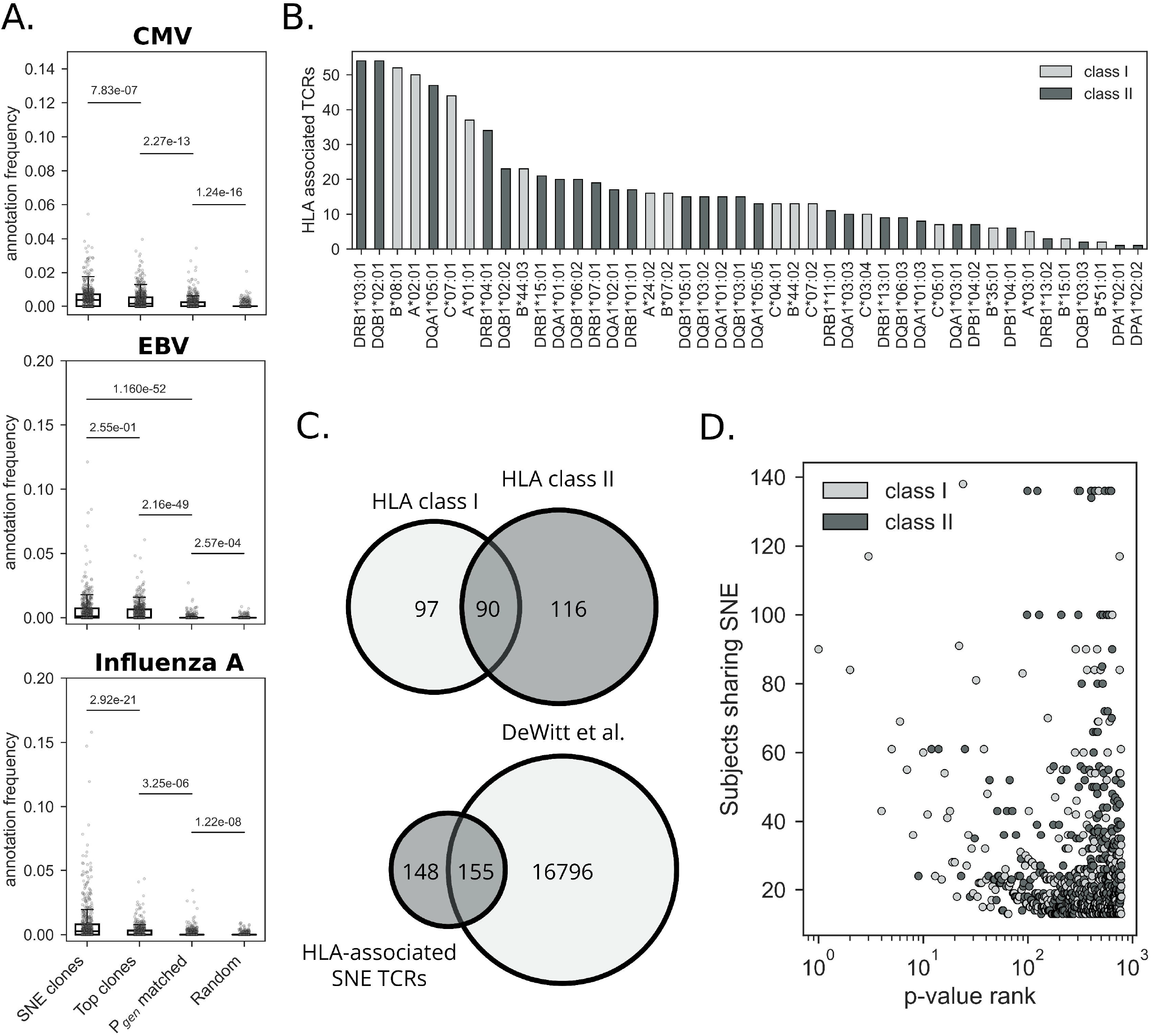
Analysis of SNE clones in the Emerson cohort. **A**. VDJdb annotation of SNE clones. 83 (out of 666) subjects were excluded from this analysis, as no SNEs were detected in these samples. For the remaining 583 subjects, we selected an equal number of the top clones (by templates). In addition, we included a Pgen matched, as well as a random sample as controls. **B**. Number of HLA-associated SNE clones per allele. B. Overlap between HLA class I- and II-associated SNE clones. **C**. Overlap between HLA class I- and II-associated SNE clones (top) and overlap of HLA-associated SNE TCRs and HLA-associated TCRs from DeWitt et al. (bottom). **D**. P-value rank of significant HLA-associated TCRs versus the number of subjects in which this TCR is neighbor-enriched. The color indicates the HLA class to which the SNE clone was associated, with the strongest associations observed for class I.

### SNE testing improves resolution of TCR-HLA associations

While no cell phenotyping information was available for the TCRs in the Emerson cohort, we can use HLA genotype of the donors to infer likely CD4 or CD8 origin of each SNE clone. First, we identified TCRs that were neighbor-enriched in >1 participant (public SNE clones). We then constructed a binary matrix with dimensions *m* x *n*, where *m* is indexed by participant and *n* is indexed by public SNE clones. We identified the top 50 HLA alleles (19 class I, 31 class II) based on public SNE prevalence in a cohort of 630 individuals and assessed the statistical significance of potential HLA-enrichment of SNE TCRs for each HLA allele using Fisher’s exact test. This resulted in the identification of 303 unique public SNE TCRs that had a positive association to at least one HLA allele (after multiple hypothesis correction, *FDR*<0.05) (**Figure 4B**). Public SNE TCRs were associated with both class I and class II HLA alleles. For 172 out of the 303 HLA-associated TCRs, we observed a significant association to >1 HLA allele, suggesting that these clones could be broadly associated to a certain haplotype due to HLA linkage disequilibrium (LD). 90 of 303 public SNE TCRs had a statistically significant association with both a class I and class II allele (**Figure 4C**), again possibly due to LD at the MHC locus. There was no difference in the number of HLA-associated TCRs between class I and class II alleles when accounting for allele frequency in the cohort (*p*=0.77, **Figure S11A**,**B**). We next compared the HLA-associated public SNE clones with the list published by DeWitt et al. (2018) [17] to compare how defining public responses based on neighbor enrichment impacts the identification of HLA-associated TCRs. While there was an overlap of 155 TCRs between the two lists, 148 of the 303 public SNE TCRs were not previously identified (**Figure 5C**). By comparing the overlapping and non-overlapping SNE clones we saw no differences in terms of the number of subjects sharing them (*p*=0.20, **Figure S11C**). However, the TCRs that were uniquely HLA-enriched among the SNE clones had higher generation probabilities, compared to the SNE clones that overlapped with the DeWitt list (*p*=2.60e-06, **Figure S11D**). Overall, HLA-associated TCRs that were SNE had higher P_gen_ (*median*=1.49e-08) compared to the HLA-associated TCRs identified through systematic enrichment testing across public clonotypes (*median*=2.87e-09, *p*=1.96e-35, Mann-Whitney U rank test). SNE testing therefore permits detection of HLA association even when publicness is partly driven by recombination bias, provided these TCRs participate in reproducible, non-random sequence neighborhood structure across individuals. Finally, despite the larger number of HLA class II associations, we found substantially more class I-associated SNE clones among the top 50 (39 class I vs 11 class II) most HLA-associated clones (**Figure 5D**).

## Discussion

Interpreting similarity in the TCR space has been a long-standing challenge. A common approach to address this challenge is through the use of similarity-based clustering methods. Performant immune receptor clustering methods are available [18–20], but these models often fail to provide meaningful interpretation to the grouped clonotypes, as they are primarily designed to group sequences into buckets of high sequence similarity without formally taking into account the biased nature of the V(D)J recombination process. This bias will result in certain clusters being more likely to emerge than others. Therefore, TCR clustering methods often miss a crucial interpretive step to account for these inherent biases, making it difficult to distinguish between clusters of TCRs that are convergently selected and those that arise purely by chance due to stochastic variation during VDJ recombination.

In the present study, we developed a framework for efficient and accurate approximation of TCRdist distances. This enabled us to evaluate the sequence neighbor density around every TCR on a repertoire-wide scale. While we believe this framework may enable development of a wide range of performant TCR analysis tools, our emphasis lay on the functional interpretation of sequence similarity enrichment in the repertoire. Specifically, to examine the hypothesis that the formation of similarity clusters is influenced by factors beyond V(D)J recombination alone (e.g., viral exposures), a background model that captures the V gene frequency, CDR3 length and generation probability distribution of the source repertoire is essential. While previous studies on sequence similarity enrichment have largely focused on single-chain data, this framework is also readily applicable to both single-chain and paired-chain αβTCR data.

In our initial validation, we found that memory T cell fractions exhibit notably higher levels of neighbor-enriched sequences in comparison to naive T cells in both mice and humans. Concurrently, no difference was observed between EM and CM fractions. Our results confirm other studies that observed greater sequence similarity in memory versus naive T cell fractions [11]. Neighbor enrichment was nearly absent among the umbilical cord blood (UCB) samples in the Britanova et al. “ages” cohort, which are known to comprise mostly naive T cells [21,12]. UCB TCRs also tend to be close to germline, with very few n-insertions due to the lower activity of terminal deoxynucleotidyl transferase (TDT) at this stage of development [22,23]. Therefore, the potential of observing many sequence neighbors among this population is high, yet controlled for by our shuffling procedure. The absence of SNEs detected in UCB is consistent with the lack of significant antigen-driven clonal expansion because the pre- and neonatal immune system has had minimal exposure to foreign antigens. In contrast, the young and middle-aged groups contained significantly more SNE clones, suggestive of more antigen exposure over time. Interestingly, neighbor enrichment was lower among the oldest individuals. In elderly individuals, the TCR repertoire often becomes oligoclonal due to repeated antigen exposure resulting in the proliferation of a few selective clones [24–26]. The proliferation of specific clones may therefore reduce the overall TCR repertoire diversity, and if these clones do not have many sequence-similar neighbors, neighbor enrichment will likely decrease. This hypothesis would align with the observations that TCR repertoire diversity decreases with age [17,27,28]. Collectively, these results suggest that our model was able to capture the baseline properties and correct for the inherent germline proximity and reduced junctional diversity seen in cord blood TCRs, ensuring that the neighbor enrichment observed in memory T cells is reflective of true antigen-driven diversification rather than innate sequence similarities driven by V(D)J recombination biases and TDT activity levels.

When applied to a longitudinal YFV vaccination data set, we hypothesized that neighbor enrichment would correspond with TCRs experiencing vaccine-induced clonal expansions. Interestingly, we found that only a small fraction of TCRβ clonotypes expanded after vaccination were also significantly neighbor-enriched. In addition, we identified many more clones that were seemingly unrelated to the vaccination stimulus, but whose convergence in the repertoire might be driven by prior or concomitant exposure to other viral epitopes. An essential finding is that neighbor enrichment analysis captured other VDJdb-annotated YFV-reactive clones that could not have been identified by looking at clonal expansion alone.

We further recognized that incorporating information from both α and β chains could improve the resolution of these enrichment analyses. This is illustrated by the fact that during the acute phase of SARS-CoV-2 infection, nearly 20% of the neighbor-enriched αβ clones could be linked to 2 known SARS-CoV-2 epitopes, after accounting for MAIT cells. In addition, we observed that numerous SARS-CoV-2-specific clones expand independently, without membership within a polyclonal response of similar TCRs (SNE cluster). This pattern likely reflects a T cell response to a broad array of epitopes, which is characteristic of responses to complex antigens. Relying solely on clonal abundance or SNE alone to monitor immune responses may therefore overlook important aspects of T cell repertoire dynamics. Sequence neighbor enrichment captures the breadth of the response at the sequence level, highlighting convergent responses that abundance metrics might miss. Incorporating this strategy alongside conventional clonal tracking may therefore enhance the assessment of vaccine responsiveness.

Finally, we observed high levels of epitope-specificity annotation for SNE TCR sequences, when compared with clonally expanded TCRs. This effect was strongest for CMV and Influenza A virus, while insignificant for EBV when matching TCRs from the Emerson cohort against VDJdb. Similarly, we observed high rates of annotation for the same viral epitopes for the individuals in the YFV cohort, where neighbor-enriched sequences were compared to longitudinally expanders. Again, higher rates of CMV epitope-specific TCRs were observed among SNE clones, along with a higher proportion of EBV epitope-specific TCRs. Coincidently, we observed that the rearrangement probabilities of SNE clones were consistently higher than those of highly abundant clones in the repertoire. This observation suggests that SNE clones may be more predisposed to recognize common viral epitopes, such as those from CMV and Influenza A. This pattern aligns with the idea that public TCRs, which have been shown to have higher rearrangement probabilities in general [29–31], could play a significant role in mounting responses to recurrent or latent viral infections [30,32]. Collectively, these results lead us to hypothesize that exposure to viral epitopes may stimulate the selection of similar clones that share specificity for these epitopes to mount an effective immune response.

### Strengths

This study presents a robust and generalizable framework for assessing TCR sequence similarity enrichment based on neighborhood density. By leveraging efficient vectorization techniques, our approach enables rapid approximation of TCRdist distances and facilitates the analysis of large-scale single- and paired-chain TCR repertoires. Importantly, our framework employs individual-specific background models which may provide more stringent neighbor-enrichment baselines than population-averaged approaches such as OLGA. This ensures that the observed neighbor enrichment accurately reflects antigen-driven selection rather than inherent biases in the V(D)J recombination process. Furthermore, the open-source nature of our tool allows for flexible integration of alternative distance metrics and provides standalone functionalities for both distance computation and background repertoire generation. This adaptability makes our framework a valuable resource for researchers exploring diverse aspects of TCR repertoire analysis.

## Limitations

Despite these promising findings, our approach has several limitations. First, we used a fixed radius of 12.5 to define TCR sequence neighborhoods in single-chain repertoire data, which may not capture all relevant TCRs, particularly in cases of rare, long rearrangements. In addition, we relied on a single distance metric, TCRdist. While various benchmarks have shown the robustness of this metric in capturing global epitope specificity, the metric may be less effective than other approaches in the context of some specific epitopes [33, 34]. Moreover, while longer vector representations (>15 dimensions per amino acid) improve the accuracy of distance approximations, they can significantly slow down computational performance, posing challenges for deeply sequenced TCR repertoires. Finally, using absolute numbers of significantly neighbor-enriched clones as a metric for sequence convergence could be misleading, as larger repertoires tend to have denser networks, which inflates the probability of observing a given number of neighbors. A more nuanced approach may be required to fully capture sequence convergence across diverse repertoires. Looking forward, the vectorization techniques employed in this study open avenues for exploring more continuous measures of TCR distance. This could enable a finer-grained analysis of neighbor enrichment, moving beyond discrete thresholds and potentially revealing subtle patterns of TCR convergence.

