## Supplementary figures for "Deciphering antigen-driven T cell responses through vectorized TCRdist sequence neighborhood quantification"

[1]Department of Computer Science, University of Antwerp, Antwerp, Belgium
[2]Antwerp Unit for Data Analysis and Computation in Immunology and Sequencing, University of Antwerp, Antwerp, Belgium
[3]Vaccine and Infectious Disease Division, Fred Hutchinson Cancer Center, Seattle, United States
[4]Translational Sciences and Therapeutics Division, Fred Hutchinson Cancer Center, Seattle, USA
[5]Division of Medical Oncology, Department of Medicine, University of Washington, Seattle, USA
[6]Public Health Sciences Division, Fred Hutchinson Research Center, Seattle, United States
[*]These authors contributed equally.

### Supplementary figures

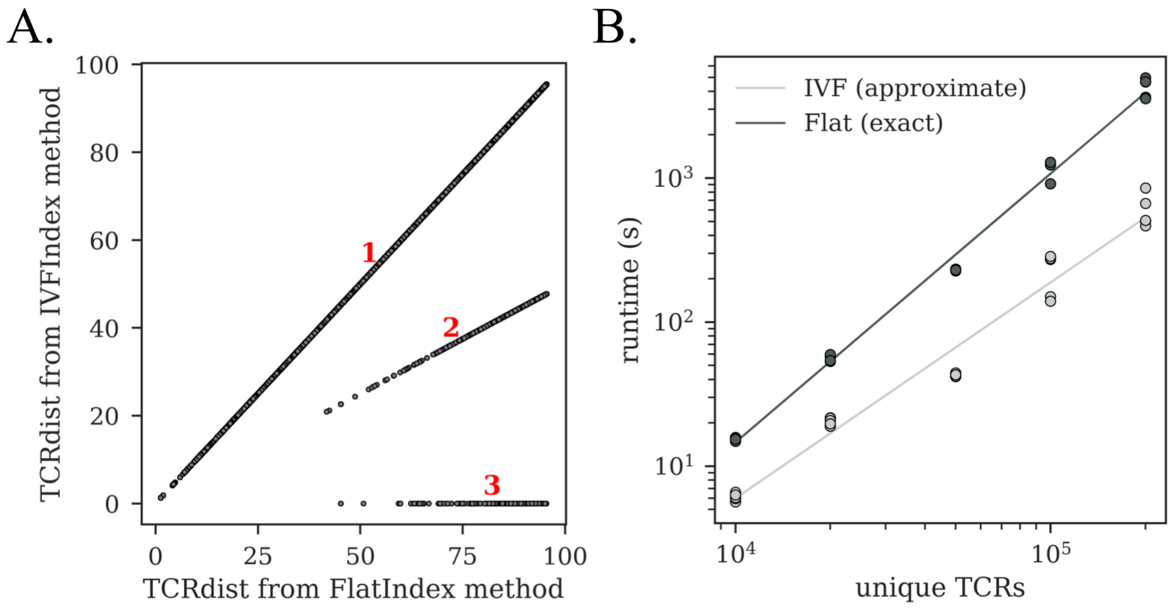


**Figure S1. Pairwise TCRdist distance calculation with clustcrdist. A.** Comparison of TCRdist distances of TCR pairs between the FlatIndex and IVFIndex methods for 200,000 unique αβ T cell clones. For this comparison, we used a 16-dimensional projection for each amino acid, and limited the maximum distance to 96.5. In total, the exact method (FlatIndex) identified 185,205 unique pairs with TCRdist < 96.5. For the large majority (183,752; 99.2%), there was agreement between both methods (1). There was another group of data points (1,087; 0.6%) for which the distance reported by the IVFIndex method was exactly half the true distance (2). Finally, only a tiny fraction of the sequence pairs (366; 0.2%) was missed by the approximate method (3). **B.** Algorithmic runtime for computing paired chain TCRdist distances across datasets with different sizes (10,000 - 200,000 TCRs).

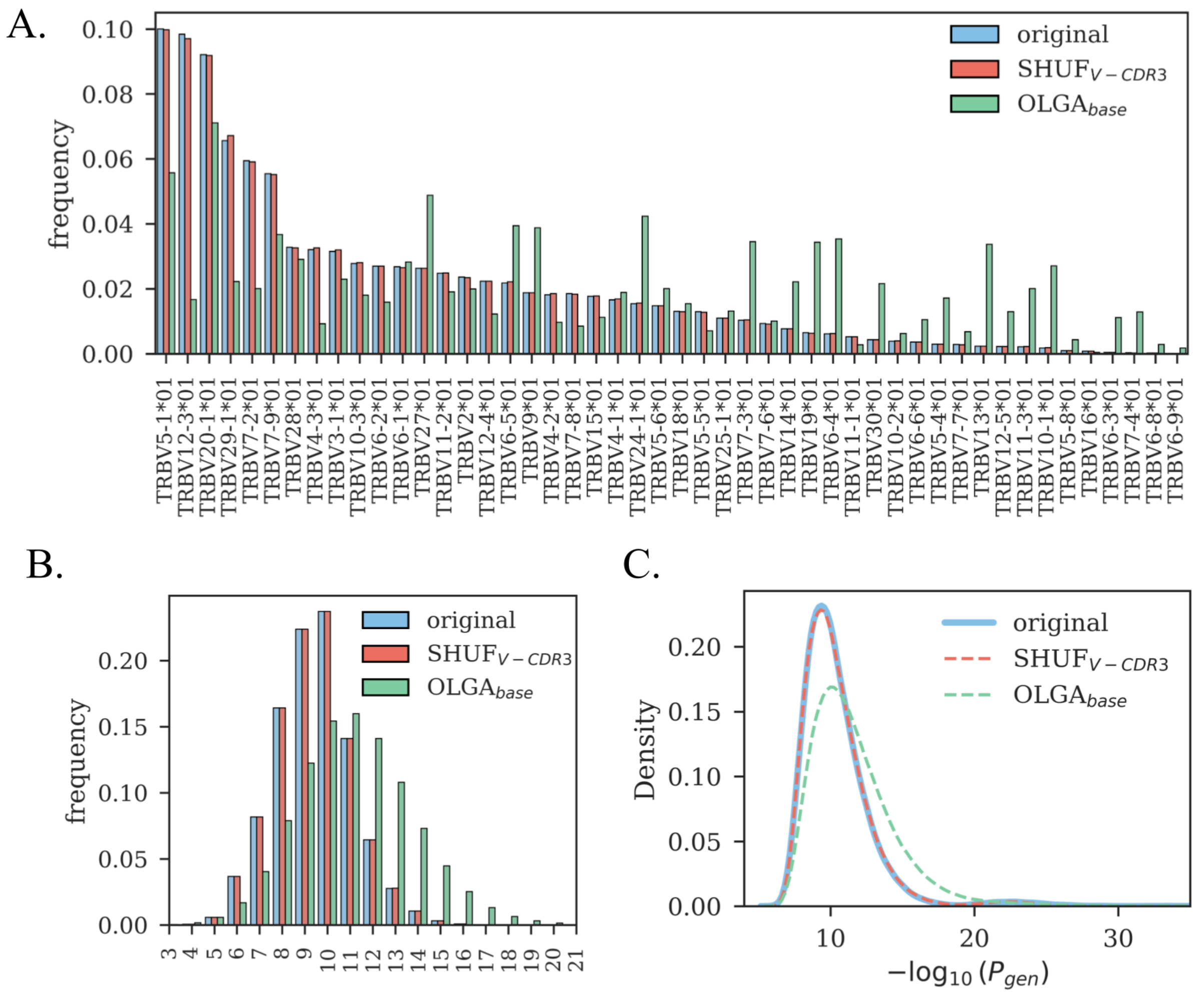


**Figure S2. Background model features. A.** V allele frequency distribution of background repertoires generated using the OLGA$_{base}$ and SHUF$_{V\text{-}CDR3}$, compared to the original input repertoire. **B.** CDR3 amino acid length distribution of background repertoires generated using the OLGA$_{base}$ and SHUF$_{V\text{-}CDR3}$, compared to the original input repertoire. **C.** Generation probability distribution of background repertoires generated using the OLGA$_{base}$ and SHUF$_{V\text{-}CDR3}$, compared to the original input repertoire.

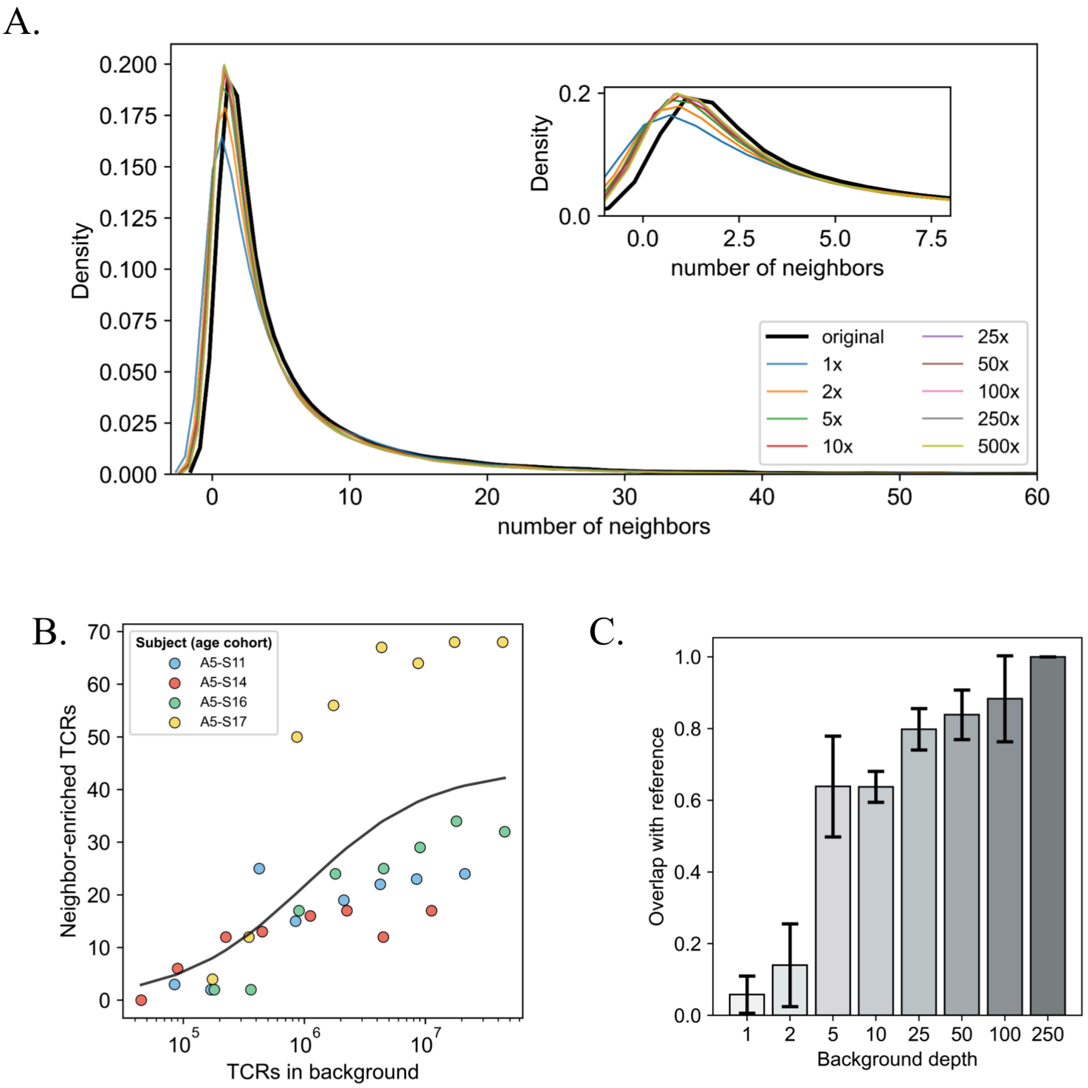


**Figure S3. Background depth. A.** The distribution of neighbors within the original repertoire (subject A5-S17; *ages cohort*) and backgrounds of differing sizes using the $\mathrm{SHUF}_{\text{V-CDR3}}$ model. Background sizes range from 1x to 500x the size of the original repertoire. The inset focuses specifically on the lower end of the distribution, spanning from 0 to 8 neighbors. **B.** Relationship between the number of TCRs in the background (generated using the $\mathrm{SHUF}_{\text{V-CDR3}}$ method) and the number of identified neighbor-enriched sequences. Each data point represents a different ratio of background TCRs to the original repertoire. The colors each represent a different input repertoire. **C**. SNE retrieval at different background sizes, ranging from 1x to 250 the size of the original repertoire. For every size, we measured the degree of overlapping SNE clones with the reference (deepest background – 250x input repertoire).

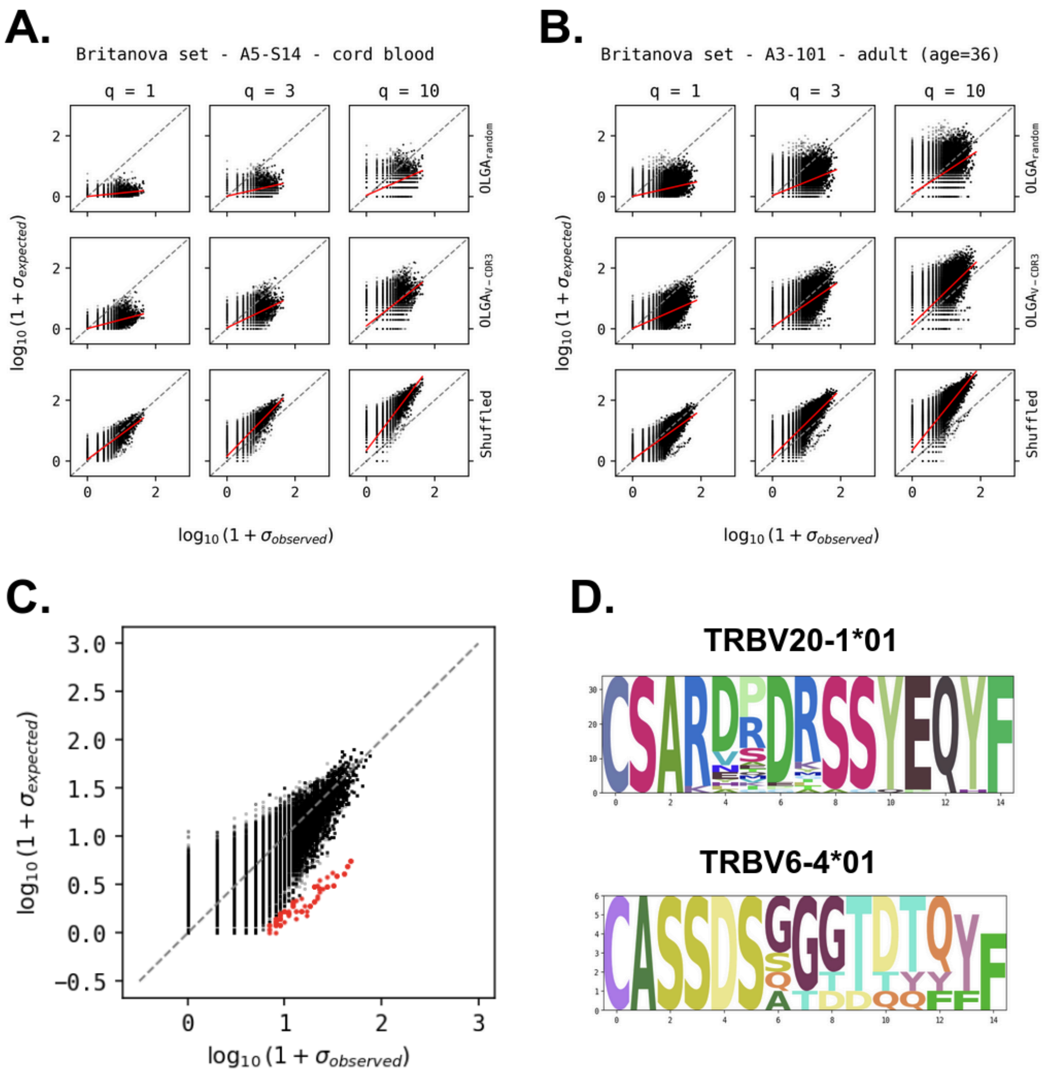


**Figure S4.** Observed versus expected neighbor counts. **A-B.** q-value correction for different background models in a cord blood (A.) and adult (B.) sample. $q$ serves a correction factor to accommodate thymic selection. OLGA models underestimate neighbor count, while the $\mathrm{SHUF}_{\text{V-CDR3}}$ preserves the neighbor distribution observed in the reference. **C.** The $\mathrm{SHUF}_{\text{V-CDR3}}$ background model reveals a population of neighbor-enriched clones (data points in red). **D.** Motifs of two neighborhoods captured by the neighbor-enriched clones in C are suggestive of MAIT TCR beta chains, which show a preference for TRBV6 and TRBV20 family V genes.

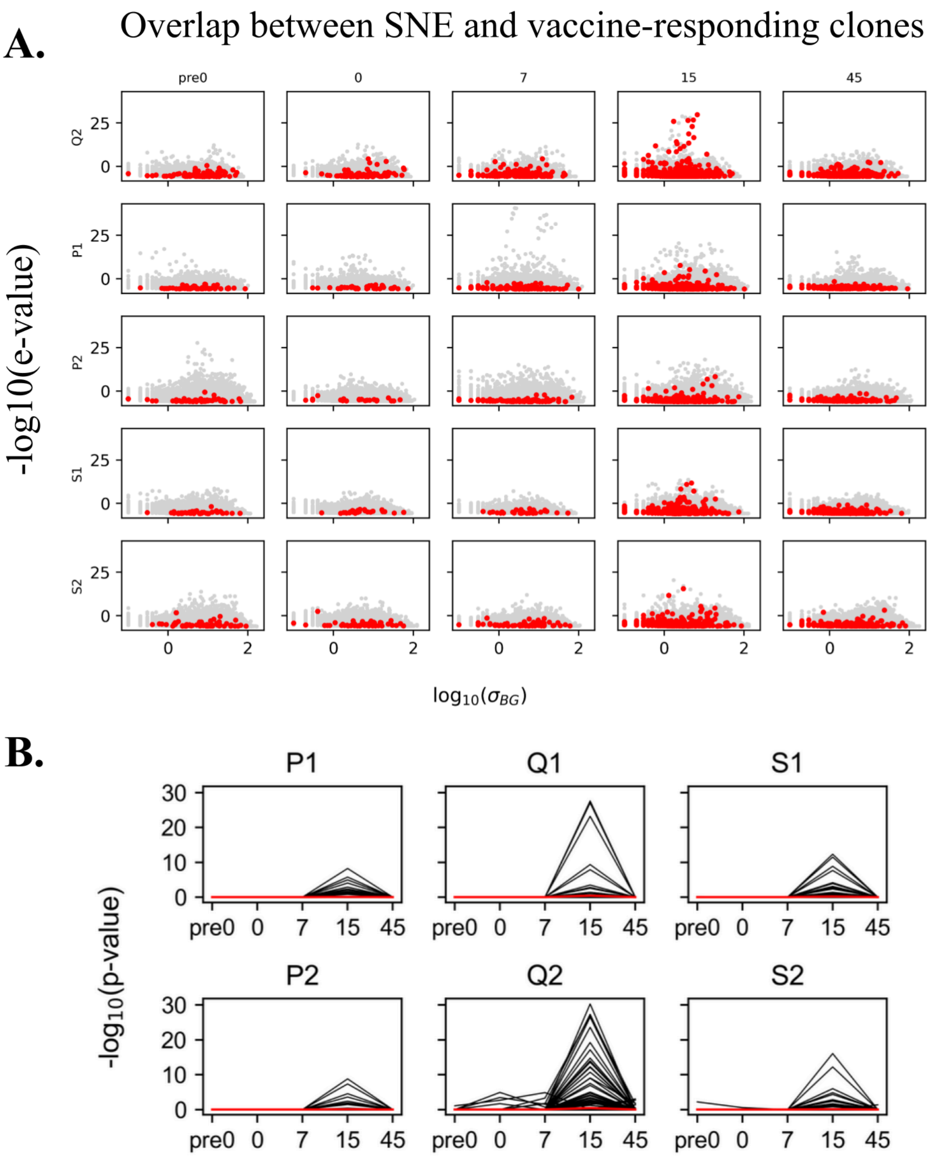


**Fig. S5. Evaluating TCR neighborhood enrichment during YFV vaccination follow-up. A.** Yellow fever virus vaccination induces convergent selection. Neighborhood enrichment was determined across 5 timepoints, for 3 pairs of twins who were administered a yellow fever virus vaccine. The 5 different timepoints included pre-vaccination, and 0, 7, 15, 45 days post-vaccination. In every participant, YFV vaccination induced convergent selection in at least a proportion of SNEs. σBG, number of background neighbors. **B.** TCR neighborhood enrichment across different timepoints during YFV vaccination. Vaccine-responding clones are exclusively neighbor-enriched at the day 15 post-vaccination timepoint. The red line shows the average -log10(p-value) for all vaccine-responding clones at each timepoint.

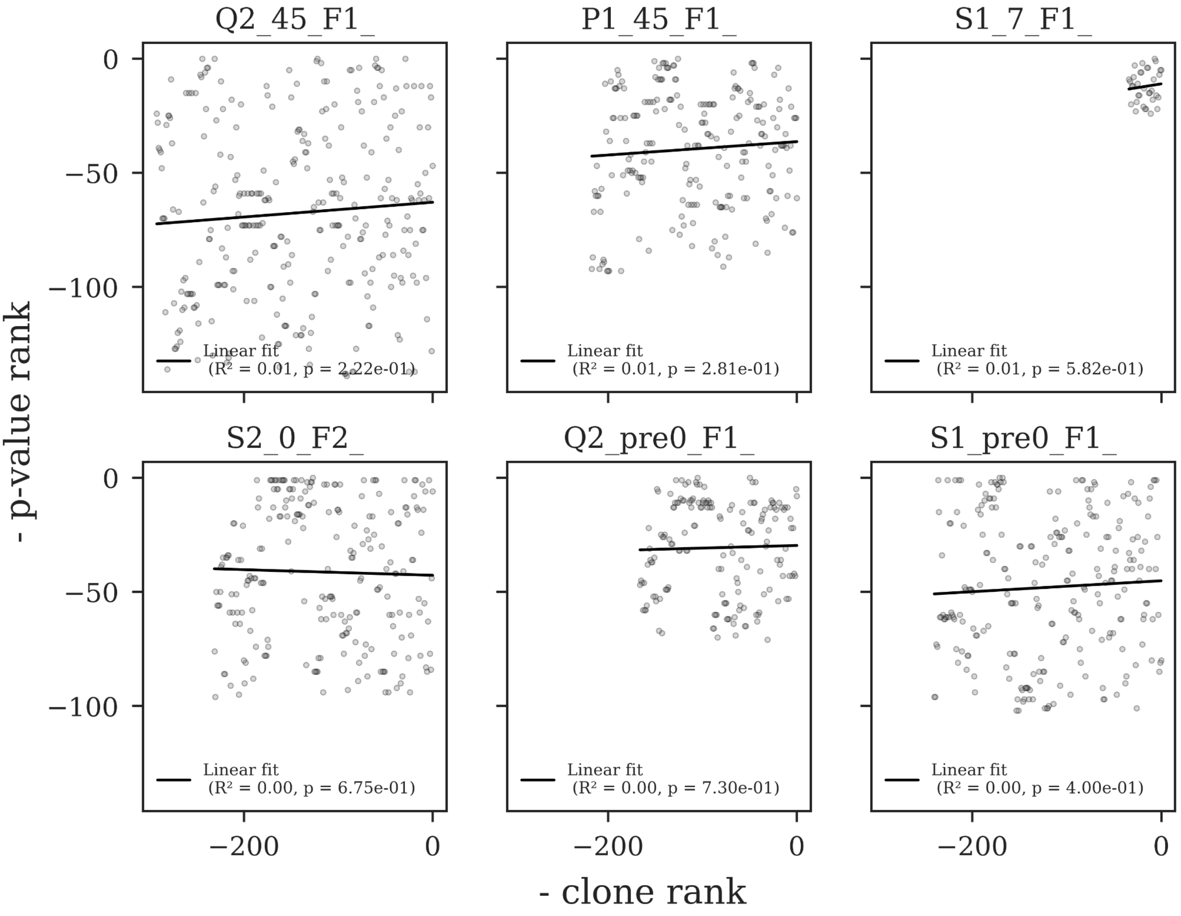


**Fig. S6. Relationship between neighbor enrichment and clonal abundance.** No correlation was found between the (negative) neighbor enrichment p-value rank and clone rank (by abundance) in any of the surveyed repertoires. Each panel represents a unique TCR repertoire.

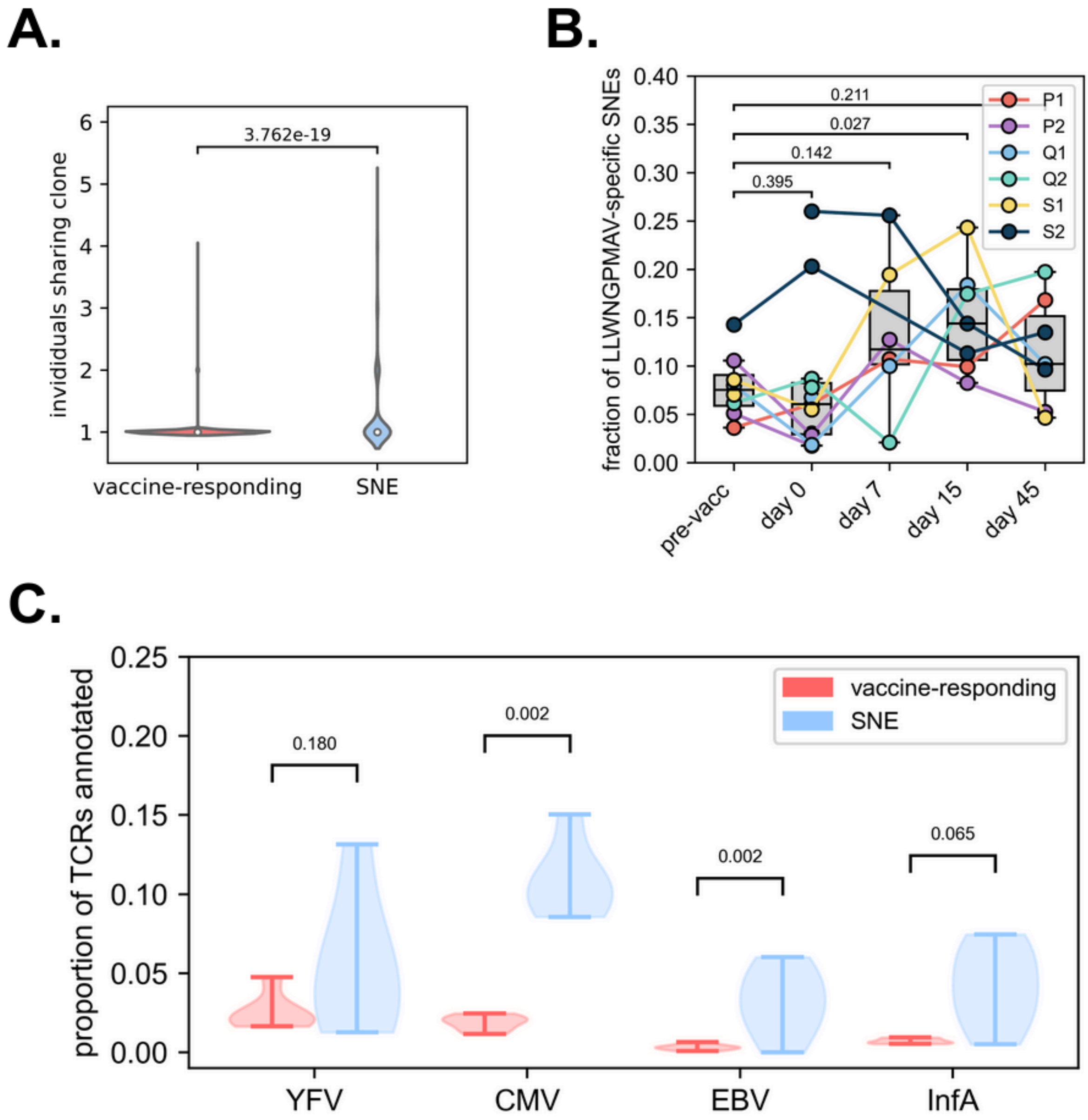


**Fig. S7.** Comparison and annotation of vaccine-responding clones and SNE clones. **A.** Number of different subjects sharing vaccine-responding or SNE clones. Stronger sharing of SNE clones is observed, relative to sharing of vaccine-responding clones (p = 3.726e-19, MWU), indicating that SNE responses are more likely to be shared among multiple individuals. **B.** Annotation of SNE clones with YFV epitope LLWNGPMAV through fuzzy matching (TCRdist < 12.5) with LLWNGPMAV-specific TCRs in VDJdb. There is a significant increase in LLWNGPMAV annotation at day 15 post vaccination (p = 0.027, MWU), compared to baseline (pre-vaccination). No differences with baseline were observed for any other timepoint. **C.** Comparison of epitope annotation in vaccine-responding and SNE clones across different pathogens. A flexible matching criterion (TCRdist < 12.5) was used to identify hits in the VDJdb. SNE TCRs matched significantly more to CMV (p = 0.002, MWU) and EBV (p = 0.002, MWU) epitope-specific TCRs in the VDJdb. No differences were observed for annotation to YFV and Influenza A virus.

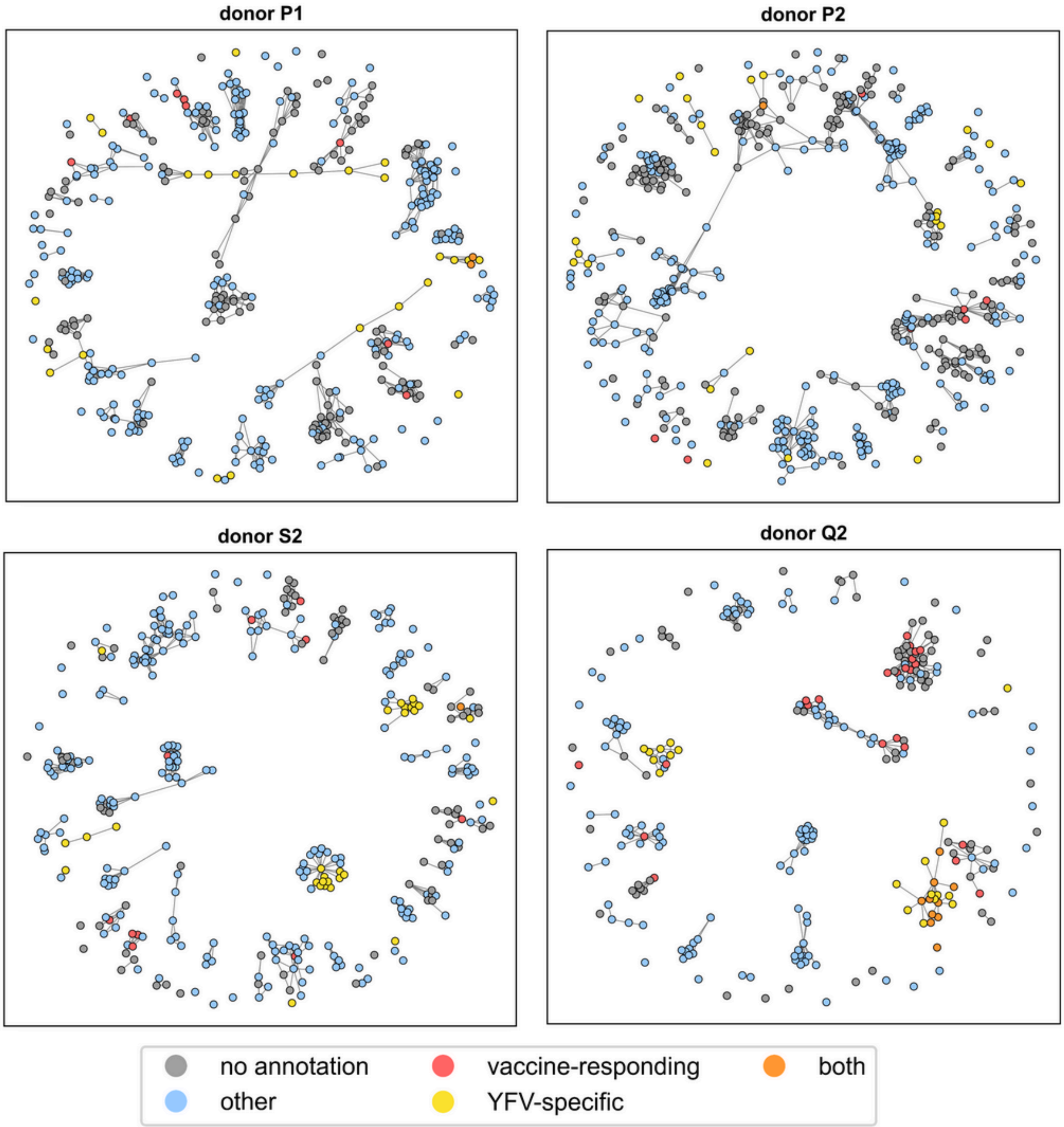


**Fig S8.** Annotation of neighbor-enriched clones at 15 days post vaccination. Red, vaccine-responding: clones that longitudinally expanded upon vaccination with the YFV vaccine; Yellow, YFV-specific: clones matching (TCRdist < 24) with a YFV epitope-specific TCR in VDJdb; Orange, both: vaccine-responding clones that also have a YFV-specific match in VDJdb. Blue, other: clones matching (TCRdist < 24) any epitope-specific TCR in VDJdb. Grey, no annotation: clones for which no annotation was found.

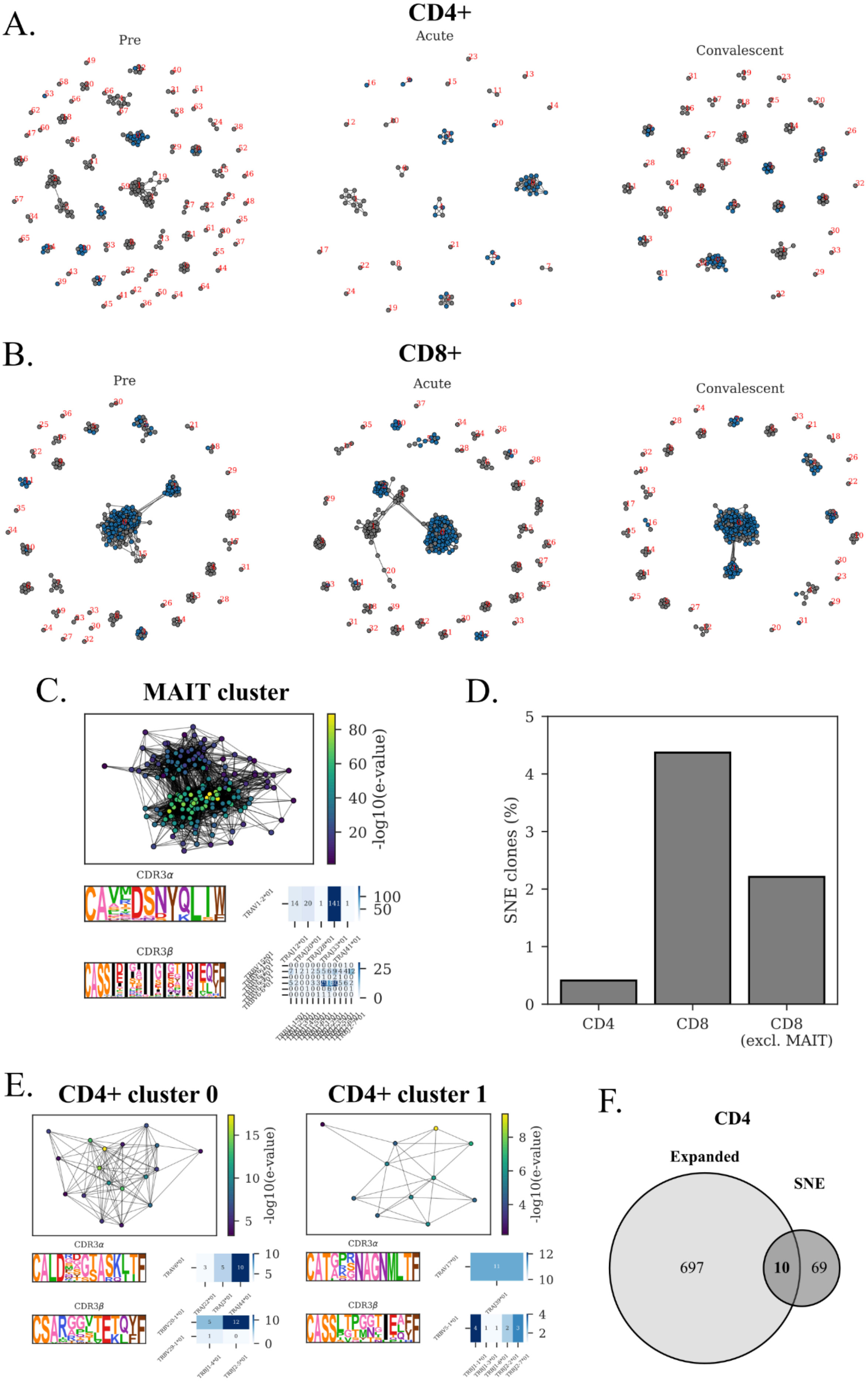
A.
CD4+
Pre
Acute
Convalescent
B.
CD8+
Pre
Acute
Convalescent
C.
MAIT cluster
-log10(e-value)
CDR3α
CDR3β
TRAV1-2*01
D.
SNE clones (%)
CD4
CD8
CD8
(excl. MAIT)
E.
CD4+ cluster 0
CD4+ cluster 1
-log10(e-value)
CDR3α
CDR3β
F.
CD4
Expanded
SNE
697
10
69

**Fig. S9. Longitudinal analysis of a single donor during SARS-CoV-2 infection.** SNE clusters at baseline (pre), acute infection and convalescence for CD4$^+$ (**A.**) and CD8$^+$ (**B.**) T cells. The sequences in blue are persistent throughout all timepoints. **C.** CD8$^+$ cluster consisting of likely MAIT cells due to the expression of TRAV1-2 and TRAJ33/12/20. The MAIT cluster is neighbor-enriched at each timepoint (cluster 0). **D.** Percentage of the TCR repertoire that is neighbor-enriched. The CD8$^+$ population contains more SNE clones (>2%) compared to the CD4$^+$ population (<0.5%). **E.** Examples of a stable SNE cluster (0, left) and a unique cluster at the acute timepoint (1, right) in the CD4$^+$ fraction. **F.** Intersection between longitudinally expanded and SNE clones among CD4$^+$ T cells.

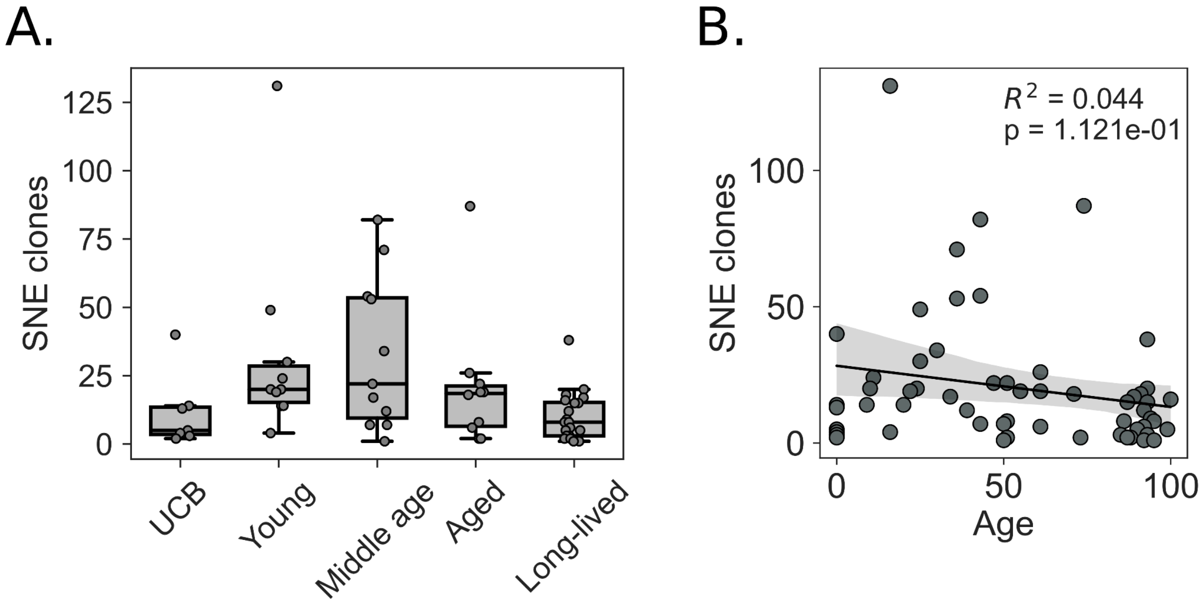


**Fig. S10. Neighbor enrichment across the lifespan. Repertoires from the *Age cohort* were downsampled to 100,000 TCRs in order to limit repertoire size bias. SNE TCR count was measured in each standardized repertoire. A.** An increase in neighbor enrichment is observed from UCB to young and middle aged individuals (p = 0.05). SNE TCR count subsequently decreased in the >= 85 years population, relative to 30 - 50 year old individuals (p = 0.04). All p-values were corrected using the Benjamini-Hochberg procedure. **B.** Linear regression analysis of SNE clone count across the lifespan, excluding UCB samples. There is a mild linear decrease in neighbor enrichment with increasing age.

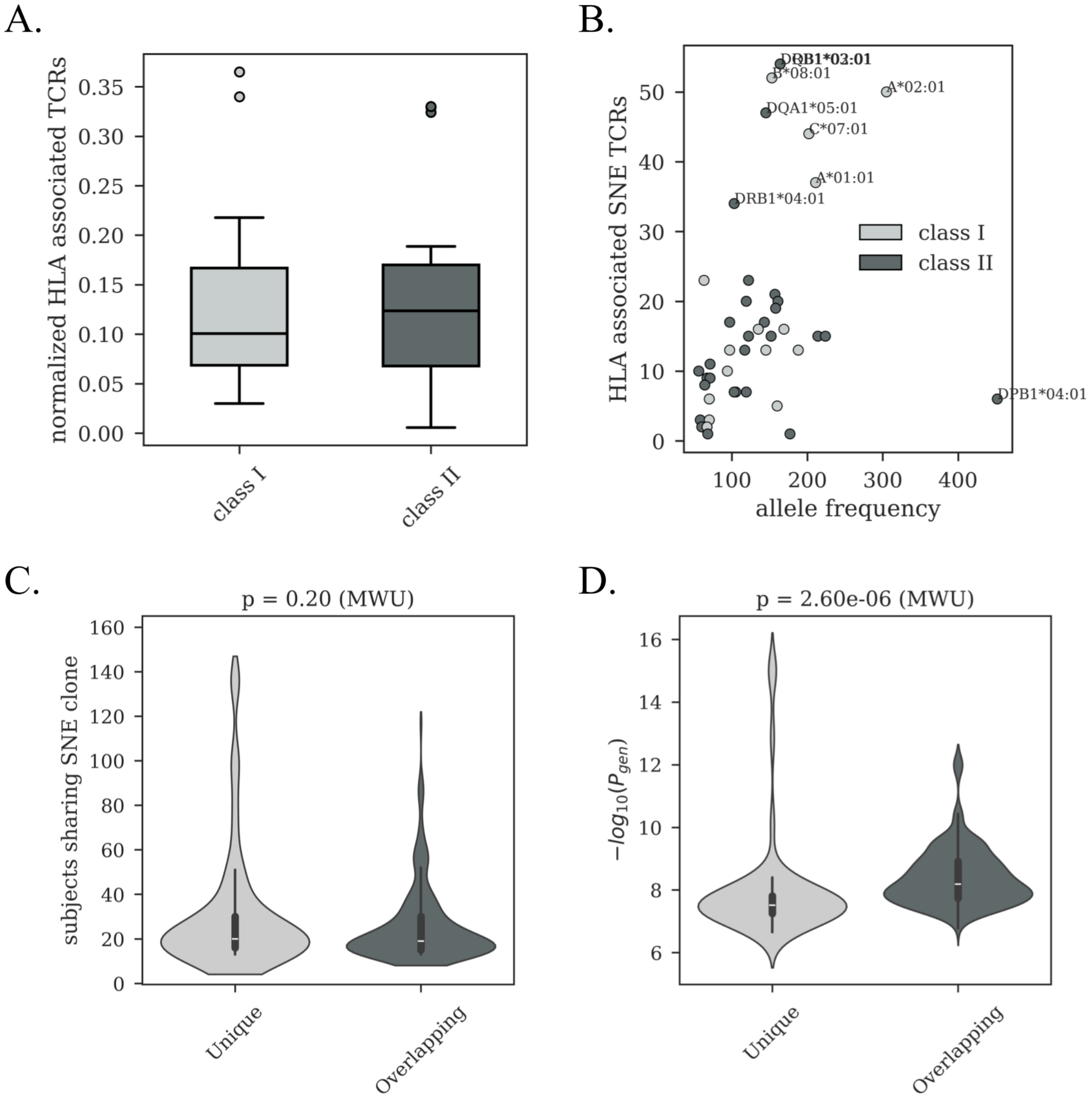


**Fig. S11. HLA association analysis of SNE clones. A.** Number of class I and II-associated SNE TCRs among subjects in the *CMV cohort*. Counts were normalized by the total number of HLA-associated SNE clones per HLA category. **B.** Allele frequency versus the number of allele-associated SNE clones for HLA class I and class II. **C.** Number of subjects sharing HLA-associated SNE, comparing clones that overlapped between SNE and DeWitt list (*Overlapping*), versus those associations that were uniquely found among the SNE clones (*Unique*). No difference was observed between the two groups (p = 0.20, Mann-Whitney U-test). **D.** -$\log_{10}(P_{gen})$ of *Unique* SNE clones versus DeWitt *Overlapping* clones. Higher $P_{gen}$ values were observed for the HLA-associated clones that were unique to the SNE list.
